# DNA Ligases Discriminate Between Natural and Non-Natural Base Pairs

**DOI:** 10.64898/2026.06.30.734188

**Authors:** Caitlin Walters-Freke, Shuichi Hoshika, Alexandra Perry, Steven A. Benner, Renwick C.J. Dobson, Zachary D. Tillett, Nigel G. J. Richards, Adele Williamson

## Abstract

Artificially Expanded Genetic Information Systems (AEGIS) increase the information content of nucleic acids by including new nucleobase pairings that are orthogonal to those of canonical Watson-Crick nucleobases. DNA ligases do not form direct interactions with the nucleobases during catalytic turnover, suggesting that these enzymes should efficiently and faithfully join double-stranded AEGIS substrates. Here we report the systematic investigation into the validity of this hypothesis for structurally-diverse DNA ligases employing substrates built from the eight nucleotide hachimoji genetic alphabet, where orthogonality is achieved by rearranging the hydrogen bonding patterns seen in canonical Watson-Crick pairs. We find that single, or multiple, non-canonical bases are well tolerated at the 5’-end of the nick. However, tracts of consecutive non-canonical bases at the 3’-end of the break significantly decrease ligation efficiency or abolish it altogether. Possible reasons for this apparent bias against non-canonical nucleobases could include incompatibility in electrostatic interactions between the ligase active site and the non-canonical substrates or altered conformational preferences and/or dynamics in key catalytic intermediates. We also observe single hachimoji mismatches are ligated more frequently than mis paired canonical bases, potentially due to promiscuous pairing of tautomeric forms of the non-canonical bases.

## INTRODUCTION

Artificially Expanded Genetic Information Systems (AEGIS) aim to generate novel nucleic acids that mimic natural DNA and RNA but have higher information density and enhanced functional group diversity (1,2). Genetic expansion is achieved by incorporating additional nucleobases that pair orthogonally to the canonical adenine-thymine/uracil (A:T(U)) and guanine-cytosine (G:C) pairs. For example, the eight nucleotide “hachimoji” (Japanese for “eight letter”) alphabet consists of the natural pairs (A:T(U) and C:G) as well as two synthetic pairs arbitrarily named **Z**:**P** and **S**:**B.** These artificial pyrimidine:purine pairs form duplexes via non-canonical arrangements of hydrogen bond donors and acceptors but still obey Watson-Crick base pairing rules, including size- and hydrogen bonding-complementarity (3).

A prerequisite for the widespread use of AEGIS alphabets in biotechnology is a suite of compatible molecular biology enzymes that permit recombination, synthesis and sequencing. DNA ligases, that can join breaks in the phosphodiester backbone of hachimoji DNA and other AEGIS nucleic acids, will be essential to generate recombinant duplexes that include non-canonical base pairs at break sites, and permit ligation-based assembly strategies for *de novo* synthesis of expanded alphabet nucleic acids (4). This latter objective has met with recent success with DNA ligase being used to install noncanonical bases including **S:B** and **P:Z** pairs into otherwise natural DNA duplexes to serve as substrates in proof-of-concept nanopore sequencing (5).

In biology, DNA ligases function to seal single strand nicks as the final step in excision repair, homologous recombination, and lagging strand DNA synthesis, or to ligate double-stranded breaks in non-homologous end joining pathways. All DNA ligases have a strict requirement for phosphorylation of the 5’ terminus of the DNA break and strongly discriminate against duplex mismatches at the junction; particularly those located at the 3’ terminus of the nick (6,7). Outside of these requirements, DNA ligases exhibit minimal preference for particular sequences in a nick ligation context although ligation rates of cohesive overhangs is influenced by G:C content in an isoform-specific manner (8). These enzymes are structurally diverse, being built around a common two-domain core comprising a nucleotidyl transferase domain (NTase domain) which carries out catalysis, and an oligonucleotide binding domain (OB domain) essential for engaging the DNA substrate (9,10). Most DNA ligases also have accessory DNA-binding domains (DB domain) or flexible loops that improve the affinity of the ligases for their nicked DNA substrates by forming a clamp-like encirclement of the duplex (11,12).

Given that ligases have evolved to join breaks in faithfully annealed DNA duplexes without significant bias towards particular sequences, we predicted that structurally diverse wild-type (WT) DNA ligases should be capable of joining the additional hachimoji base pairs **S:B** and **Z:P** with a similar efficiency and fidelity to their canonical (A:T, C:G) counterparts. Structural and computational studies have shown that non-canonical pairs in hachimoji DNA retain the major features of natural DNA, such as an equivalent duplex diameter, major and minor groove widths lying within the range observed of G:C and A:T pairs, and the ability to interconvert between A- and B-forms in a manner similar to that of natural DNA (13–15). These properties suggest that the ‘sequence agnostic’ behaviour of DNA ligases will extend to non-canonical nucleobase pairs, such as those comprising the hachimoji alphabet, thereby positioning these enzymes as ideal tools for assembly of AEGIS nucleic acids.

Here we report a systematic study of AEGIS DNA ligation that examines the generalisability of non-canonical nucleotide joining by a structurally diverse group of ligases. We examine the efficiency of ligating non-canonical base combinations at the 5’- and 3’-termini of a nick, as well as the effect of consecutive non-canonical bases at the break. We then investigate the fidelity of AEGIS ligation by examining equivalent substrates which present a mismatch against canonical bases.

From an enzymology perspective, interrogation of the ligase reaction with various non-canonical bases that deviate from natural nucleotides in defined ways provides insights into the limits of DNA ligase catalysis and the molecular determinants of end recognition. From an engineering perspective, although we have developed optimal reaction conditions for ligating substrates containing tracts of consecutive non-canonical hachimoji nucleotides we also show that this is at the further expense of ligation fidelity.

## MATERIALS AND METHODS

### Recombinant expression of DNA ligases

Recombinant DNA ligases from *Alteromonas mediterranea* (Ame-Lig), *Burkholderia pseudomallei* (Bsp-Lig), *Rhizobium phage* vB_RleM_P10VF (P10VF-Lig) and *Acinetobacter phage Ac42* (Ac42-Lig) as well as *Prochlorococcus marinus* ligase LigP (Pmar-LigP) and LigW (Pmar-LigW) were expressed and purified as described previously (16–19). Details of the DNA ligases used in this study are given in Supplementary section 1. For all recombinant DNA ligases, the clarified lysate was supplemented with 0.1 mM ATP, 10 mM MgCl_2_ and incubated at 4 °C overnight to ensure post step 1 enzyme-adenylate form was obtained.

### Synthesis of oligonucleotides and preparation of AEGIS-containing substrates

Oligonucleotides containing 2-amino-5-methyl-1-(1’-beta-D-2’-deoxyribofuranosyl)-4(1*H*)-pyrimidinone (**S**), 6-amino-9-(1’-beta-D-2’-deoxyribofuranosyl)-(1*H*)-purin-2-one (**B**) and adenine (A), guanine (G), cytosine (C) and thymine (T) were purchased from IDT. Oligonucleotides containing 2-amino-8-(1-beta-D-2-deoxyribofuranosyl)imidazo[1,2-a]-1,3,5-triazin-[8H]-4-one (**P**), 6-amino-3-(2-deoxyribofuranosyl)-5-nitro1H-pyridin-2-one (**Z**), 3-methyl-6-amino-5-(1’-β-D-2’-deoxyribofuranosyl)-pyrimidin-2-one (2’-deoxy-1-methylpseudocytidine) (**pS**) and **B** were synthesised as described previously (3).

The reason two versions **S** and **pS** are used is that initial oligonucleotides for S:B containing substrates were purchased from IDT for expediency; however, due to their synthetic protocols the S base could not be installed at the 3’ end. For in-house synthesis of oligonucleotides with this base at the 3’ end, we used the pS version as this is more stable during chemical synthesis steps.

For the oligonucleotides synthesised within this project, standard dA, dT, dG, dC phosphoramidites, d**B** (dmf-isodG-CE), 5’-Fluorescein phosphoramidite, and Chemical Phosphorylation Reagent were purchased from Glen Research (Sterling, VA). AEGIS phosphoramidites (d**Z**, d**P**, and d**pS**) were purchased from Firebird Biomolecular Sciences LLC (Alachua, FL). All oligonucleotides containing AEGIS nucleosides were synthesized on an ABI 394 DNA Synthesizer using standard phosphoramidite chemistry. The controlled pore glass supports (CPGs) carrying oligonucleotides were treated with 2.0 mL of 1 M DBU in anhydrous acetonitrile at room temperature for 24 h to deprotect the *p*-nitrophenylethyl group on the d**Z** nucleobase. Then the CPGs were filtered, dried, and treated with concentrated ammonium hydroxide at 55°C for 16 h. After removal of ammonium hydroxide, the hachimoji oligonucleotides were purified on 20% denatured PAGE and then desalted using Sep-Pac® Plus C18 cartridges (Waters). Structures of the AEGIS bases are shown in Figure 1 A and B. Hachimoji and regular oligonucleotides were resuspended and annealed as described previously (20) to make a range of substrate duplexes containing ligatable nicks shown in Figure 1 C. The individual oligonucleotide sequences and the combinations used to produce the various hachimoji substrates are given in Supplementary sections 2A and 2B.

**Figure 1.**
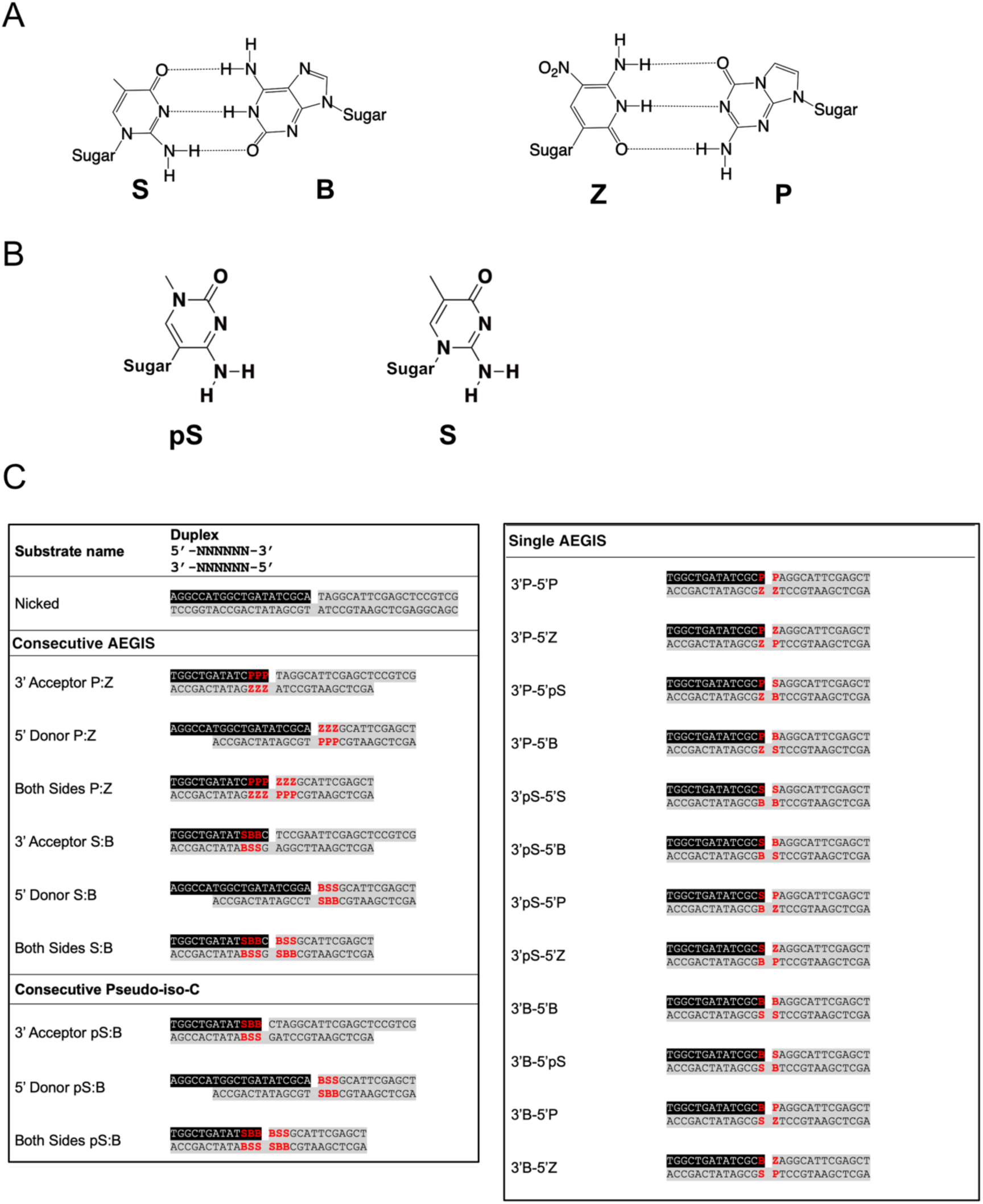
A) Structures and base pairing interactions between the non-canonical nucleotides in 8-nucleotide hachimoji DNA. B) Structure of iso C (2-amino-5-methyl-1-(1’-beta-D-2’-deoxyribofuranosyl)-4(1H)-pyrimidinone); hereafter **S**); and pseudo-iso C, (3-methyl-6-amino-5-(1’-β-D-2’-deoxyribofuranosyl)-pyrimidin-2-one (2’-deoxy-1-methylpseudocytidine); hereafter **pS**). Note that **S** and **pS** present identical hydrogen bond donor-acceptor patterns and both interact with **B** as shown in the panel above. C) Hachimoji and standard DNA duplexes used in this study. Black shading indicates 5’ 6-fluorescein labelled strand used for detection; red text indicates AEGIS bases. The upper strand is written 5’ à 3’, the complement is presented in the 3’à 5’ orientation for clarity. The individual oligonucleotide sequences and combinations annealed to generate these are provided in Supplementary sections 2A and 2B.

### Hachimoji DNA ligation reactions

The ability of various DNA ligases to join substrates containing different combinations of hachimoji DNA nucleotides was measured using the fluorescently-labeled duplexes given in Figure 1 C and resolved by denaturing urea PAGE as described previously (17,20). Standard reaction components (80 nM double stranded DNA oligonucleotides, 1 mM ATP, 10 mM Mg^2+^, 50 mM TRIS pH 8.0, 50 mM NaCl, 1 mM DTT) were used to evaluate the relative activities of the various ligase:substrate combinations with incubation for 30 min at 25 °C. The impact of temperature (15 °C) and time (30 min – 18 h) on Pmar-LigP and Ac42-Lig were also determined by varying those parameters using the standard reaction components. Steady-state reaction kinetics were determined using the standard reaction component mixture but with varying concentrations of natural or hachimoji DNA substrate (20nM – 300 nM) and with enzyme concentrations of 10 nM (natural DNA), 20 nM (3’ acceptor S:B and 3’ acceptor P:Z) for Pmar-LigP and 1nM (natural DNA), 20 nM (3’ acceptor S:B and 3’ acceptor P:Z) for Ac42-Lig.

‘Optimized’ reaction conditions for enhancing hachimoji ligation were identified by varying components of the standard reaction mix including the following changes: the crowding assay involved addition of 10% w/v crowding agents, which included either PEG 3350, PEG 6000, PEG 8000, betaine or glycerol. The ATP-concentration assay involved varying the concentrations of ATP from 0.1 mM to 5 mM. The impact of metal ion concentration and type was assessed by substituting the 10 mM Mg^2+^ with either Mg^2+^ or Mn^2+^ at concentrations between 0.5 mM and 50 mM. Where Mn^2+^ was used, the pH of the buffer was lowered to pH 7.0 to avoid precipitation. For all assays, reaction products were resolved by electrophoresis on 20% denaturing TBE urea gels (7 M urea, 20% acrylamide-Bis 29:1, 1x TBE) and visualized on an iBright imager (Invitrogen) using the fluorescein setting. Image quantification was done using ImageJ (https://imagej.nih.gov/ij/). Urea-PAGE gel images are available on Zenodo (DOI:10.5281/zenodo.20821393).

### Hachimoji DNA binding measured by microscale thermophoresis, analytical ultracentrifugation and electrophoretic mobility shift assay

All binding assays were carried out on the same batches of enzyme which were pre-treated with ATP/MgCl_2_ during purification as described above, to ensure a consistent adenylation state.

Microscale thermophoresis (MST) measurements were carried out as described previously (18). Substrates were prepared as described for EMSA binding assays but with the addition of 0.05% tween20. DNA substrate was incubated with 0.2–35 µM of each DNA ligase for 15 min at 4 °C. MST measurements were carried out on a Nanotemper Monolith NT.115 using standard capillaries and data was analyzed using the MO.Affinity software.

Sedimentation velocity type analytical ultracentrifugation (SV-AUC) experiments were performed with a Beckman Coulter Optima analytical ultracentrifuge using an AN-50 Ti 8-hole rotor. Sample (380 μL) and buffer (400 μL) was loaded into cells with a 12 mm epon 2-channel centerpiece and either quartz or sapphire windows. For the data shown in Figure 5D, Pmar-LigP was serially diluted to a range of final concentrations (0.18–6.2 μM). The final concentration for the FAM-labelled hachimoji substrates was ∼1 μM. The buffer composition of the protein-DNA complex was 50 mM Tris pH 8.0, 75 mM NaCl, 1 mM DTT, 1 mM EDTA, 2.5% v/v glycerol. Data were collected at 20 °C and at 50,000 rpm, acquired in absorbance mode at 495 nm (exploiting the absorbance of the FAM-label on the hachimoji DNA), using radial absorbance scans at a step size of 0.001 cm. Samples were incubated at on ice for at least 3 hours, followed by a further 1.5 hours at room temperature as the sedimentation experiment was setup and the rotor came to temperature (20 °C) prior to starting the experiment to ensure the Pmar-LigP and hachimoji substrate mixtures approached equilibrium and to minimize protein aggregation. To standardize the experiments (which have different ratios of hachimoji DNA to protein), the partial specific volume (v-bar) is 0.73 mL/g for the macromolecules, buffer density of 1.00 g/mL, and viscosity of 1.002 cp. Sedimentation velocity data were fitted to a continuous *c(s)* distribution model using the program SEDFIT (21,22). In these experiments we track the hachimoji DNA and the fraction bound to protein is determined by integrating the continuous *c(s)* distribution using GUSSI (version 2.1.0, (23). The dissociation constant (*K*_d_) was determined using the standard quadratic equation.

Hachimoji DNA binding was measured using electrophoretic mobility shift assay (EMSA) on native PAGE as described previously (18,20). Briefly, the binding assay mixture included the same components as the standard ligation reactions however the 10 mM Mg^2+^ was substituted with 5 mM EDTA. Binding was initiated by the addition of DNA ligase enzyme at 0.8 μM final concentration and samples incubated at 15 °C for 30 minutes. After this time, loading buffer was added (100 mM EDTA, 0.25% bromophenol blue, 25% v/v glycerol) and samples were electrophoresed on 10% native TBE gels (10% acrylamide-Bis 29:1, 1x TBE) and visualized on an iBright imager (Invitrogen) using the fluorescein setting.

## RESULTS

### Diverse DNA ligases can join duplexes with consecutive AEGIS base pairs

To determine how effectively DNA ligases can join sequential non-canonical nucleotides in hachimoji DNA, and to assess the impact of ligase structure on joining efficiency, we determined the activity of six structurally-divergent ligases on duplexes containing tracts of consecutive AEGIS bases (Figure 2 A and B). All DNA ligases tested are ATP-dependent and include the ‘minimal’ two-domain ligases (Bsp-Lig and Ame-Lig), three enzymes with alpha helical ‘T4-like’ DNA binding domains (Pmar-LigP, P10VF-Lig and Ac42-Lig) as well as one ligase with an unusual predicted beta sheet DNA-binding domain (Pmar-LigW).

**Figure 2.**
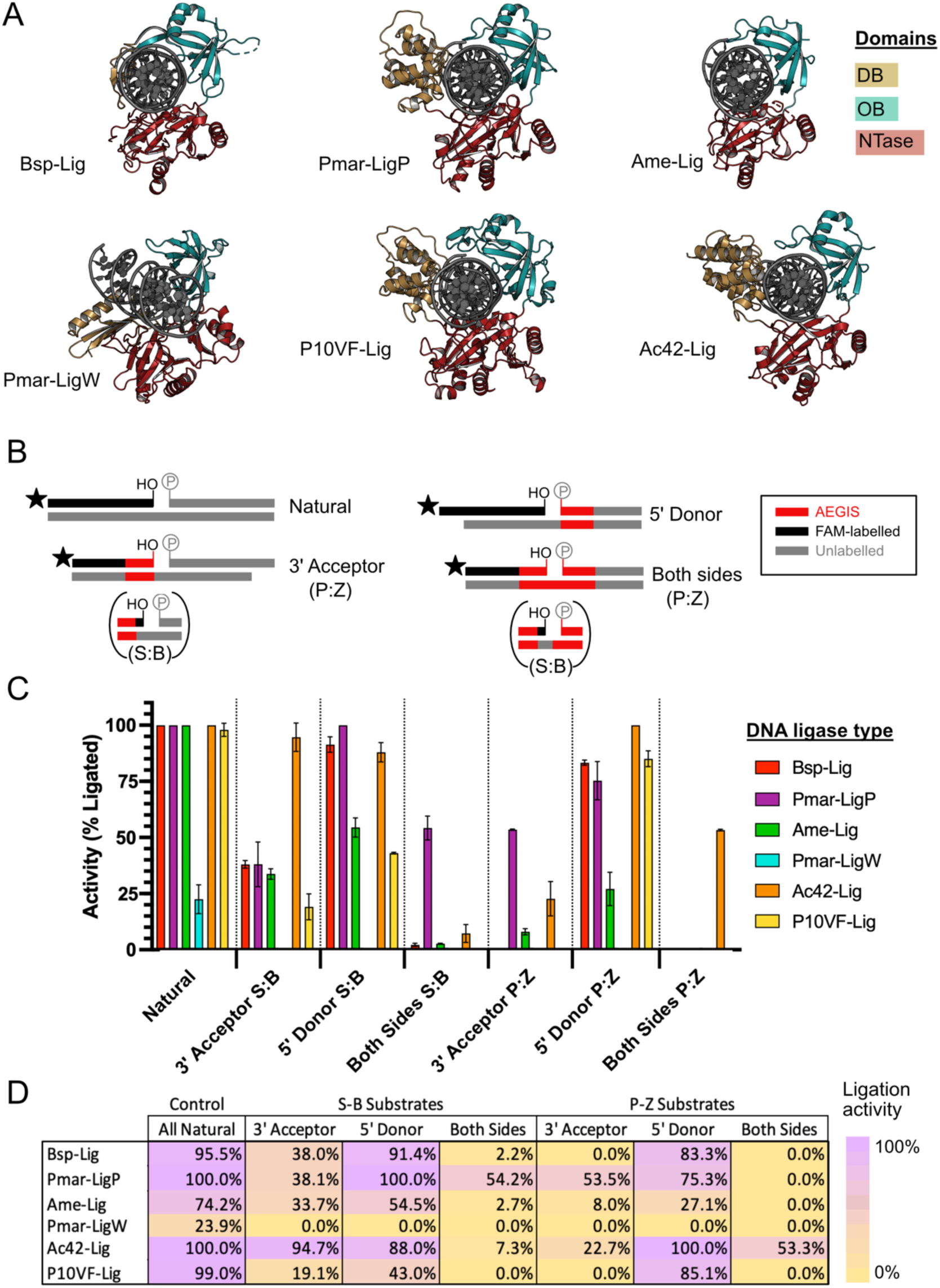
A) Structures of DNA ligases used in this study. Bsp-Lig, 7OBN; Pmar-LigP, 6RAR; Ame-Lig, 6GDR; P10VF-Lig, 9FZX. Pmar-LigW and Ac42-Lig were modelled using AlphaFold3 bound to nicked DNA (43). B) Schematic of hachimoji substrates containing consecutive **S:B** or **P:Z** pairs. Note that due to synthesis limitations, commercial oligonucleotides containing **S** and **B** used for initial experiments had a natural nucleotide (cytosine) in the 3’ terminal position. Consequently, direct **S:B** to **S:B** ligation was not tested with this initial series. C) Extent of ligation by different DNA ligases quantified from urea-PAGE analysis of fluorescently labelled substrates (raw gel images are given in Supplementary Figure 3). The final enzyme concentration was 56 nM for Pmar-LigW and 20 nM for all other ligases tested. Incubation times were 30 min at 25 °C. Values are the mean of three replicates for 3’ acceptor **S:B** substrates and two replicates for others; error bars are the standard error of the mean. D) Comparison of the effect of AEGIS base position on ligation summarised as a heat map. Values are equivalent to Panel C.

The most significant difference in activity between substrates is seen when multiple non-canonical nucleotides are present in the acceptor strand at the 3’ end of the nick (Figure 2 C and D). A run of three **S**:**B** pairs reduces the percentage ligated to below 40% for almost all enzymes tested, while a run of three **P**:**Z** pairs reduces the percentage ligated to below 10% for most enzymes. Inclusion of three consecutive AEGIS bases on the 5’ donor strand has a relatively minor effect on activity for most of the ligases tested when these are in the context of natural nucleobases on the 3’ acceptor end. However, the presence of non-standard pairs on both sides of the break is extremely deleterious to ligase activity, essentially abolishing joining in the case of **P**:**Z** pairs, and decreasing it below 10% for **S**:**B** pairs.

The extent of activity also depends on the specific ligase enzyme used. In general, the helical DB-domain enzymes produce the highest yields of joining for all hachimoji substrates with both Ac42-lig and Pmar-LigP having the best joining ability of substrates with AEGIS bases in any position. The two minimal ligases tested, Bsp-Lig and Ame-Lig have overall poorer activity with Ame-Lig being particularly impacted by AEGIS bases on the 5’ donor strand, relative to the other ligases. The unusual Pmar-LigW ligase produces no discernible activity on any of the hachimoj substrates; however this enzyme also exhibits poor activity on the control nicked substrate suggesting that this may reflect an overall low rate of DNA ligation rather than a bias against inclusion of non-canonical bases in the duplex.

To determine whether the presence of a natural nucleobase at the 3’ terminus influences the efficiency with which multiple non-standard pairs could be ligated, a new set of oligonucleotides was synthesised in-house where **B** was at the 3’-end of the DNA ligation substrate. These were tested with the two most efficient ligases, Ac42-lig and Pmar-LigP. The C-nucleoside **pS** version was used in place of **S** this new oligonucleotide series, as this form is less prone to deamination (24); for this reason, ligation of substrates incorporating **pS** at equivalent positions as the **S** base in the 5’-donor strand were also directly compared.

Introduction of the three consecutive **pS:B** pairs directly in the 3’ acceptor strand shows no detectable joining of either the substrate with **pS:B** bases on the 3’ acceptor strand, or the substrate with **pS:B** on both sides of the nick (Figure 3). In contrast, little difference in ligation yield is seen when comparing the 5’ donor substrate with three consecutive **pS:B** bases and the original with **S:B** pairs. Taken together, this suggests that it is the presence of a consecutive run of non-standard nucleotides on the 3’ end of the ligatable nick, rather than the replacement of **S** with **pS**, which prevents effective ligation in this oligonucleotide series.

**Figure 3.**
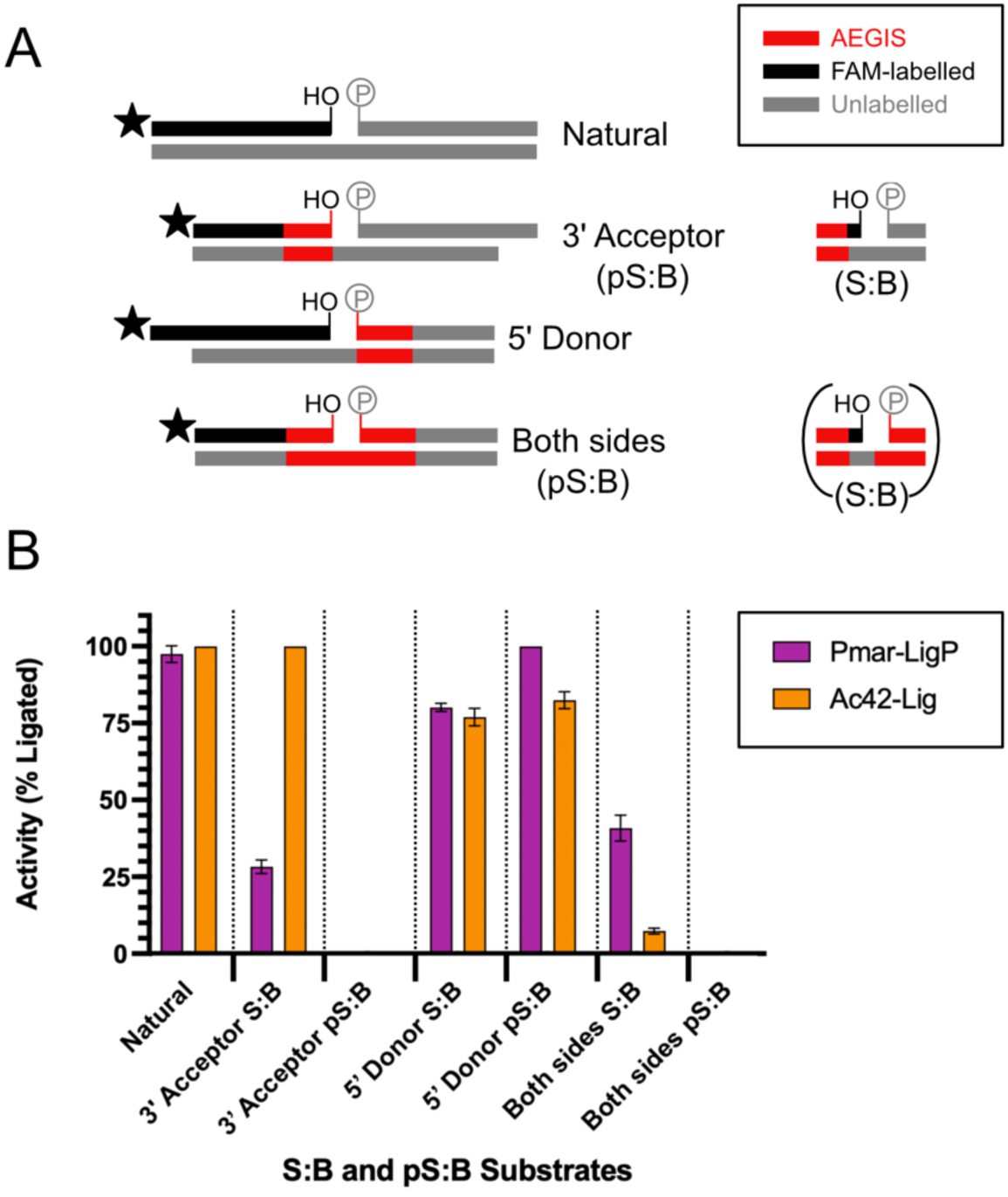
A) Schematic of hachimoji substrates used to compare the impact of the nature of the 3’ nucleotide on ligation efficiency. See Figure 1 and Supplementary 2 for substrate sequences and details. B) Extent of ligation by Pmar-LigP and Ac42-Lig when the 3’ terminal base pair is canonical or an pS-B pair. Values are the mean of three replicates; error bars are the standard error of the mean. Both ligases were assayed at a concentration of 20 nM, incubation times were 30 min at 25 °C.

### Nature of the non-standard nucleotide at either terminus influences ligation

Given that the presence of consecutive AEGIS bases on either terminus of a ligatable break impacts ligation efficiency, we wished to determine the effect of different single AEGIS bases at each side of the nick. To test this, a series of substrates were synthesised that included all combinations of **pS**:**B** and **P**:**Z** pairs at the 3’ or 5’ end of the break, except for a 3’ **Z**, which could not be incorporated in this position due to synthetic limitations. As with the previous experiment, activity was examined with the two most active ligases, Ac42-lig and Pmar-LigP (Figure 3).

The extent of ligation activity depends strongly on the nature of the terminal nucleotides (Figure 4). The combinations 3’**P**-5’**P**, 3’**P**-5’**Z**, 3’**P**-5’**pS** and 3’**P**-5’**B** give almost 100% yields with both enzymes. In contrast, the combinations 3’**pS**-5’**pS** and 3’**pS**-5’**B** are ligated at less than 40% yield by either enzyme.

**Figure 4.**
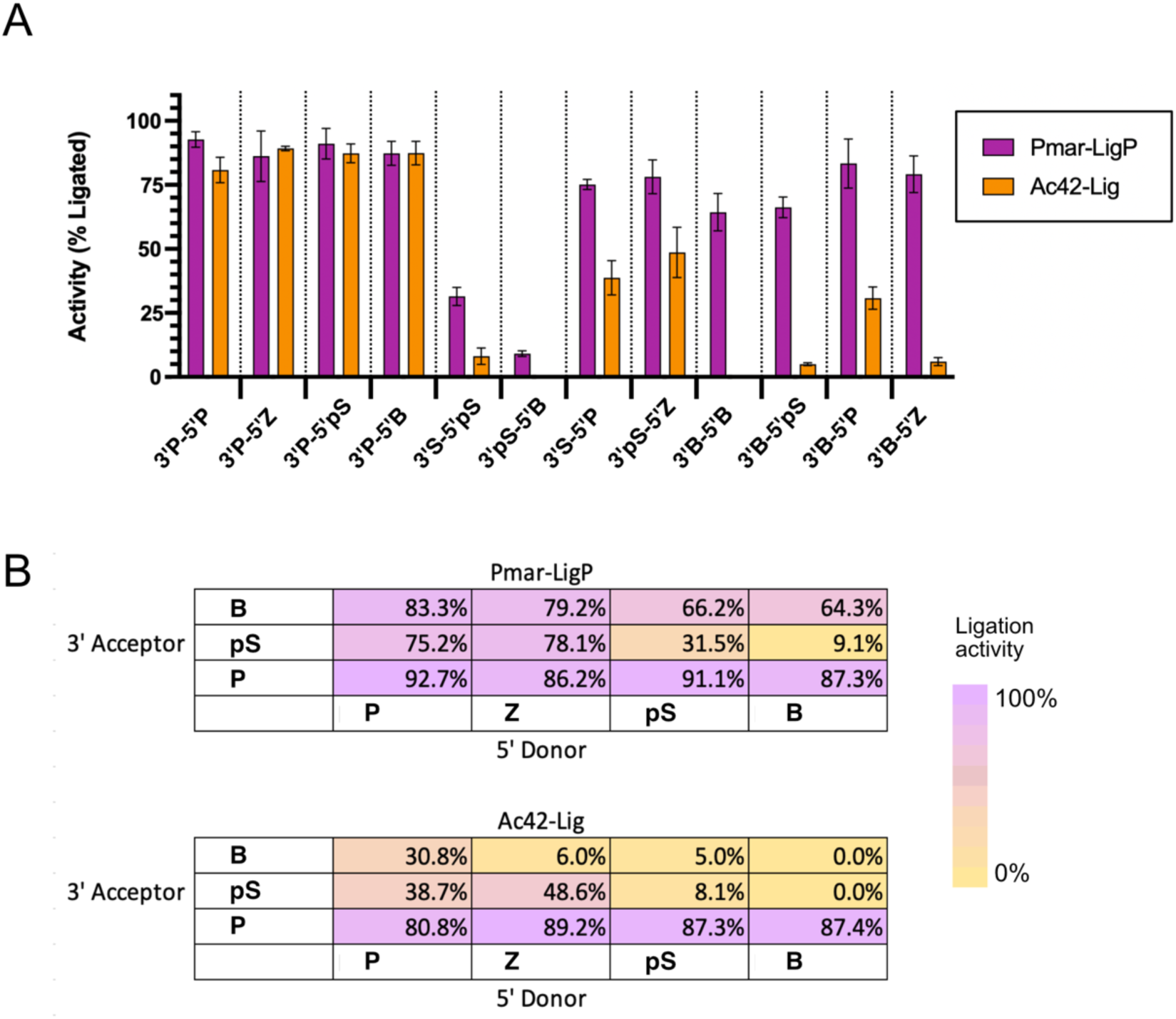
A) Ligation of single AEGIS substrates by Pmar-LigP and Ac42-lig. See Figure 1 and Supplementary 2 for substrate sequences and details. Values are from quantification of urea-PAGE analysis (Supplementary 5), and both ligases were assayed at a concentration of 20 nM, incubation times were 30 min at 25 °C. Values are the mean of three replicates; error bars are the standard error of the mean. B) Comparison of the impact of non-standard nucleotides as donor and acceptor termini represented as a heat graph of the percentage of ligation. Values are equivalent to Panel (A).

**Figure 5.**
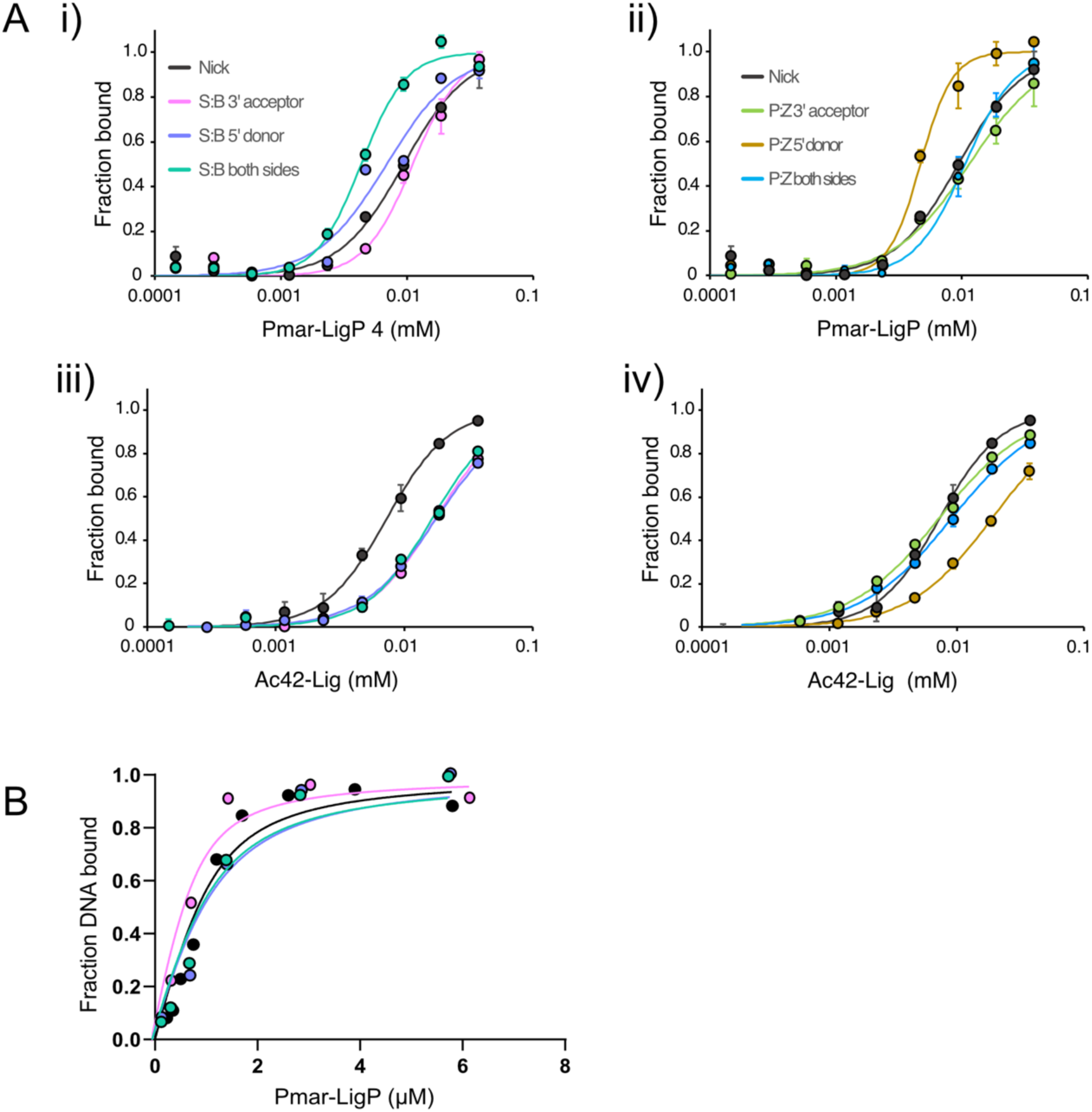
Binding of Pmar-LigP and Ac42-lig to hachimoji substrates. A) Binding evaluated by MST to substrates with consecutive AEGIS bases. Values are the mean of three replicate measurements; error is the standard error of the mean. B) Binding of Pmar-LigP to hachimoji substrates with consecutive non-standard pairs evaluated by sedimentation velocity type analytical ultracentrifugation (SV-AUC) experiments. The R^2^ for the fits are: 0.93 for natural DNA; 0.91 for S:B (3’ acceptor); 0.93 for S:B (5’ donor); and 0.94 for S:B (both sides). The distributions for each experiment from which the fraction bound was determined are included in Supplementary Figure 3, as are representative data and fits.

Compared with the results of experiments ligating nicks with multiple consecutive AEGIS bases, this indicates that both enzymes can join terminal combinations with a single **P** base at the 3’ acceptor terminus very efficiently when it is flanked by natural bases. However, in the context of adjacent non-standard pairs, such as the ligation with **PPP**-**ZZZ** pairs of the ‘**P:Z** both-sides’ substrate, efficiency is reduced. Interestingly, enzyme-specific trends vary between the two ligases. Ac42-lig shows a striking decrease in activity when a **B** nucleotide is present on the 3’ acceptor end; for example, 3’**B**-5’**B**, 3’**B**-5’**pS** and 3’**B**-5’**Z** are joined with less than 10% efficiency, while Pmar-LigP has more than 50% ligation with three.

### Defective binding of hachimoji DNA does not explain all poor ligation

To determine whether defective binding accounts for the low level of ligation activity with consecutive **S**:**B** and **P**:**Z** hachimoji substrates we measured affinity of Ac42-lig and Pmar-LigP for them using microscale thermophoresis (MST) (Figure 5A). We observe minimal difference in ligase affinity between the natural DNA and these hachimoji substrates (Table 1), and in particular there is negligible decrease in affinity of Pmar-LigP for **P:Z** (both sides) despite no ligation activity being observed for this ligase-substrate combination. Similarly, relative to natural DNA, Ac42-Lig has only a marginal decrease in affinity for the **S:B** (both sides) substrate measured by MST, despite ligating it at less-than 10%. The binding of Pmar-LigP, to consecutive **S**:**B** substrates was further verified by sedimentation velocity type analytical ultracentrifugation experiments (SV-AUC, Figure 5B and Supplementary 3) which, consistent with the MST results, shows that multiple AEGIS bases at nick termini do not significantly disrupt ligase binding. Differences in the absolute dissociation constants obtained using MST and SV-AUC are likely due to differences in the experimental setup (e.g. addition of 0.05% tween20 for protein stabilisation in MST and different incubation times); however both techniques indicate that binding is not affected.

**Table 1.** Binding affinities of ligases to hachimoji substrates determined by MST (Pmar-LigP and AC42-Lig) and AUC (Pmar-LigP). Values for MST represent the mean of three replicate measurements and error is given as the confidence value of the fitted model. For the AUC experiments, 95% confidence intervals are reported. Representative sedimentation velocity type analytical ultracentrifugation (SV-AUC) data and fits, along with the c(s) distributions from which the fraction bound is determined, are shown in Supplementary Figure 7.

| Substrate | Pmar-LigP |  |  | Ac42-lig |  |
| --- | --- | --- | --- | --- | --- |
|  | Activity (%)<sup>*</sup> | MST EC50 (μM) | AUC K<sub>d</sub> (μM) | Activity (%)<sup>*</sup> | MST EC50 (μM) |
| **Natural** |  |  |  |  |  |
| Nick | 100 | 9.4 (±2.3) | 0.34 (0.16–0.60) | 100 | 7.3 (±0.08) |
| Linear (no nick) |  | 21.13 (±1.25) |  |  | nd |
| **Multiple AEGIS bases** |  |  |  |  |  |
| S:B (3' acceptor) | 38 (±10.0) | 11.1 (±1.40) | 0.21 (0.05–0.53) | 94 (±6.3) | 17.4 (±3.30) |
| S:B (5' donor) | 100 | 6.9 (±1.50) | 0.50 (0.17–1.13) | 88 (±4.3) | 17.7 (±1.80) |
| S:B (both sides) | 54 (±5.3) | 4.7 (±0.40) | 0.44 (0.16–0.94) | 7 (±4.0) | 16.5 (±3.40) |
| P:Z (3' acceptor) | 2 (±0.1) | 11.5 (±2.00) |  | 22 (±7.7) | 7.2 (±1.10) |
| P:Z (5' donor) | 75 (±8.5) | 4.7 (±2.90) |  | 100 | 18.9 (±10.33) |
| P:Z (both sides) | 0 | 10.6 (±1.80) |  | 53 (±0.4) | 9.0 (±1.10) |
| **Single AEGIS bases** |  |  |  |  |  |
| 3'B-5'B | 64 (±7.3) | 11.8 (±0.46) |  | 0 | nd |
| 3'B-5'pS | 66 (±4.0) | 21.41 (±4.11) |  | 5 (±0.6) | nd |
| 3'B-5'P | 92.7 (±2.8) | 18.73 (±2.12) |  | 31 (±4.4) | nd |
| 3'B-5'Z | 79.2 (±7.1) | 14.18 (±2.09) |  | 6 (±1.6) | nd |
| 3'pS-5'B | 9 (±0.0) | 10.8 (±0.53) |  | 0 | nd |
| 3'pS-5'pS | 31 (±3.5) | 18.1 (±1.88) |  | 8 (±3.2) | nd |

Using MST, we then investigated ligase affinity for single AEGIS terminal combinations which were poorly ligated (Table 1). Binding of 3’B-5’B, 3’B-5’Z and 3’pS-5’B by Pmar-LigP is relatively unaffected, despite the latter being ligated very poorly, however Pmar-LigP affinity for 3’B-5’pS, 3’B-5’P and 3’pS-5’pS is decreased and has a similar affinity to the linear un-nicked control DNA (21.1 µM). By contrast, none of the poorly-ligated single-terminus AEGIS substrates are bound by Ac42-Lig to any appreciable extent when measured by MST, although qualitative EMSA assays did detect some interaction (Supplementary 4). Ac42-Lig also has no measurable binding to the linear DNA control suggesting higher specificity for ligatable nicks.

To further elucidate the cause of low ligation activity with consecutive AEGIS bases, we measured the steady-state kinetics of Pmar-LigP and Ac42-lig with multiple S:B or P:Z bases at the 3’ acceptor end. Due to substrate inhibition at DNA concentrations above 300 nM we were unable to reach saturating substrate concentrations in these assays meaning kinetic constants could not be directly derived by fitting the Michaelis Menten equation (Supplementary 5). However by substituting the EC_50_ values determined from MST (Table 1) for K_m_, we were able to define an upper limit for K_cat_ for each substrate from the rate vs [DNA] curves where the linear portion is equivalent to K_cat_/K_m_. This indicates that the catalytic step is 10-20 fold slower for ligation of termini with consecutive 3’ AEGIS bases (Table 2.).

**Table 2.** Estimated upper limit of K_cat_. Steady-state kinetics plots are provided in Supplementary figure 5.

| Protein | Substrate | Upper limit K<sub>cat</sub> (s<sup>-1</sup>) |
| --- | --- | --- |
| **Pmar-LigP** | Nicked DNA | 2.27 |
|  | S:B (3' acceptor) | 0.11 |
|  | P:Z (3' acceptor) | 0.17 |
| **Ac42-Lig** | Nicked DNA | 23.05 |
|  | S:B (3' acceptor) | 2.11 |
|  | P:Z (3' acceptor) | 0.08 |

### Addition of crowding agents enhances hachimoji DNA joining

To identify conditions that improve the ligation of hachimoji substrates, we examined the impact of varying different reaction parameters on the ligation yield when sequential non-standard pairs (**S:B** both sides and **P:Z** both sides) were joined by Pmar-Lig and Ac42-lig (Figure 6).

**Figure 6.**
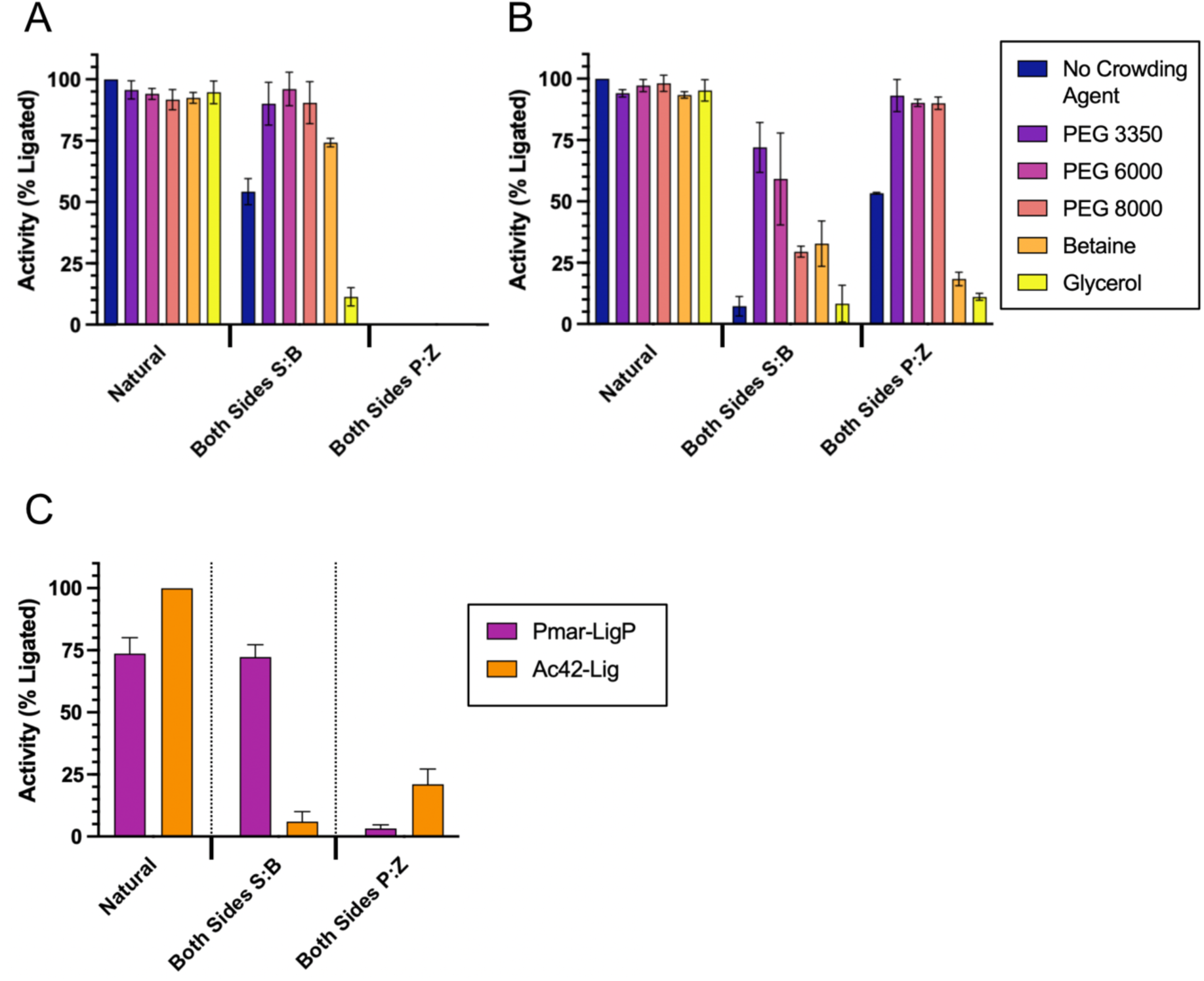
Effect of crowding agents on ligation of sequential non-standard pairs (both sides). A) Ligase activity of Pmar-LigP, incubation time 30 min at 25 °C. B) Ligase activity of Ac42-Lig, incubation time 30 min at 25 °C. C) Ligase activity of Pmar-LigP and Ac42-Lig after incubation at 15 °C overnight (no addition of crowding agents). Values are the mean of three replicates from quantification of urea-PAGE analysis error bars are the standard error of the mean. Both ligases were assayed at a concentration of 20 nM for all experiments.

Addition of crowding agents proved the most effective individual variable increasing yields, with all three PEG additives promoting ligation. The effect of PEG3350 is particularly notable in increasing Ac42-lig activity with **S**:**B** pairs (both sides) from almost negligible levels (7.3%) to over 70%, while ligation yields from sequential **P**:**Z** pairs (both sides) are also enhanced (Figure 6B). By contrast, although PEG addition stimulates Pmar-LigP joining of sequential **S**:**B** pairs (both sides) (from 54.2% to 90.0%), no detectable ligation of sequential **P**:**Z** pairs (both sides) can be observed (Figure 6A).

No improvement in ligation activity was detected with changes in ATP or MgCl_2_ concentration, or by substitution for Mn^2+^ as the divalent metal ion cofactor (Supplementary 6-9). These parameters were therefore not changed from those originally used in all subsequent experiments. Increased incubation times (up to 18 h) and decreased temperatures (15 °C) have a small but detectable impact on ligation yields. Specifically, under these conditions, ligation to form sequential **P**:**Z** pairs (both sides) by Pmar-LigP is observed, while ligation to form sequential **S**:**B** pairs (both sides) increases to over 70%, on-par with the ligation observed for natural DNA under these conditions (Figure 6C).

When combined with addition of 10% PEG3350 and a molar excess of enzyme, both Pmar-LigP and Ac42-lig can join all single nick combinations tested to more than 80% completeness. This includes consecutive **S**:**B** and **P**:**Z** nucleotides on either or both sides of the nick (Figure 7A), as well as 3’**B**-5’**P**, 3’**B**-5’**Z**, 3’**B**-5’**S**, 3’**S**-5’**S** and 3’**S**-5’**B** which AC42-Lig has especially poor activity with under standard conditions (i.e. no additives, 20 nM enzyme and incubation for 30 min at 25 °C) (Figure 7B).

**Figure 7.**
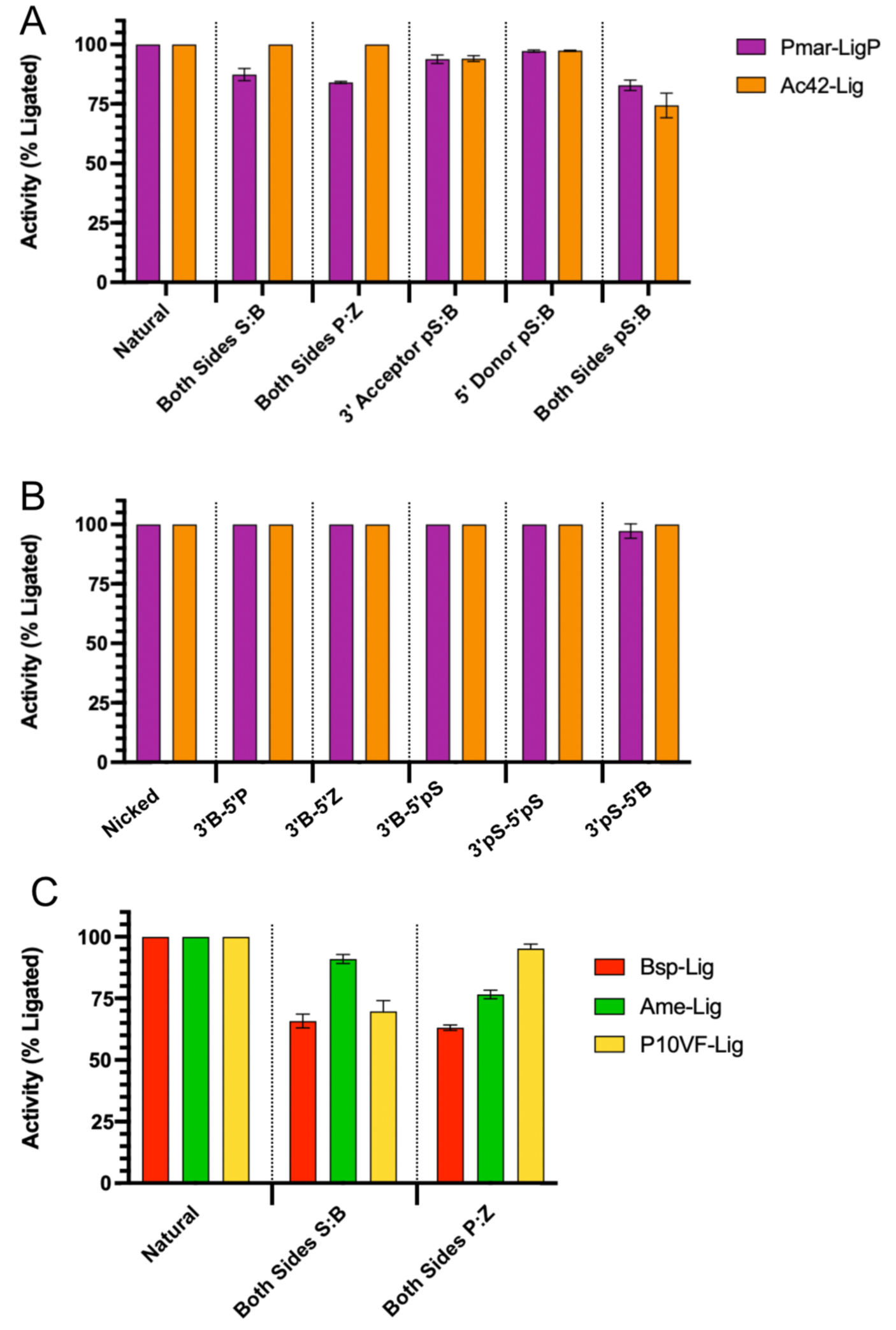
Ligation of consecutive AEGIS substrates using optimised reaction conditions: 10% PEG 3350, 18 hours incubation at 15 °C. Values are the mean of three replicates from quantification of urea-PAGE analysis and error is the standard error of the mean. Ligases were assayed at a concentration of 1.9μM for Pmar-LigP samples, 1.9 μM for Ac42-Lig samples, 3.7 μM for Bsp-Lig, 1.6 μM for Ame-Lig and 1.7 μM for P10VF-Lig.

To determine the general applicability of these conditions to increase the yields of ligation products containing consecutive non-standard base pars, we assayed three less-active DNA ligases from our original panel, Bsp-Lig, Ame-Lig and P10VF-Lig. In all cases more than 60% of substrate that contain **S**:**B** pairs (both-sides) and **P**:**Z** pairs (both-sides) substrates are ligated (Figure 7C), a considerable increase from less than 3% under non-optimised conditions.

### AEGIS bases are ligated with low fidelity and crowding agents further lower discrimination

Consistent with many characterised DNA ligases (7,25), Pmar-LigP and Ac42-Lig have low fidelity with mismatches at the 5’ end and readily join a single mis paired bases under the standard conditions used for initial experiments (20 nM enzyme, 30 min, 25°C, no crowding agent), but have much lower tolerance for 3’ mismatches with less than 25% ligation for an A:C (purine:pyrimidine) and less than 5% for the larger A:G (purine:purine) mismatches (Figure 8 Bi). However, using the ‘optimised’ reaction conditions which promote ligation of AEGIS base termini (10% PEG3350, 18 h incubation at 15 °C and molar excess of enzyme) both 5’ and 3’ mismatches are joined with more than 70% yield (Figure 8 Bii).

**Figure 8.**
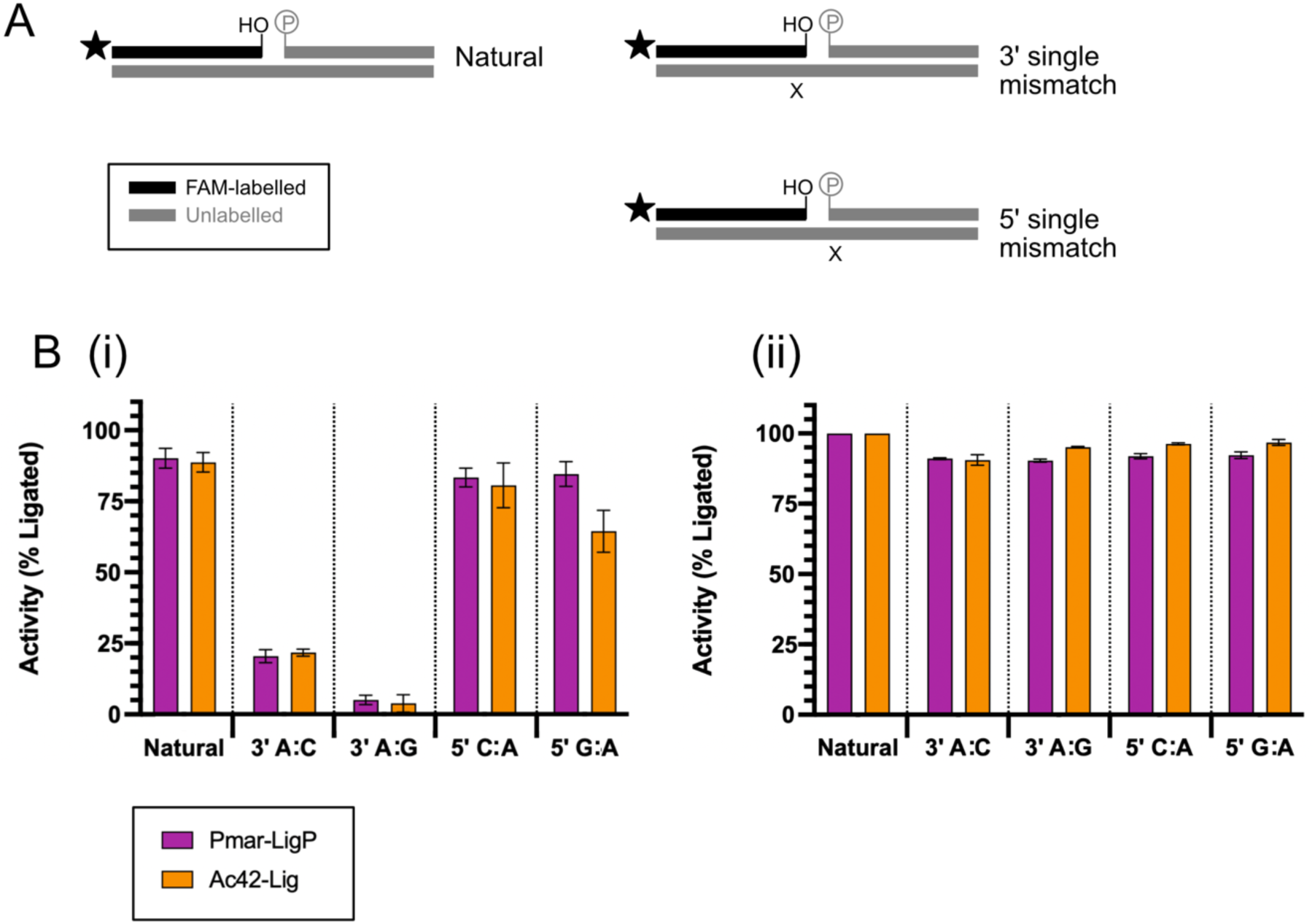
Ligation of mismatched canonical nucleobases by Pmar-LigP and Ac42-Lig. A) Schematic of DNA substrates containing mismatched natural base pairs with ‘X’ indicating the approximate position of the mismatch Oligonucleotides and duplex sequences are given in supplementary sections 2 and 10 respectively. B) Ligation yield with single mismatched bases at either terminus under standard assay conditions (20 nM enzyme and incubation time 30 min at 25 °C) (i) and optimised conditions (10% PEG 3350, 1.9μM enzyme and incubation time 18 hours at 15 °C) (ii). Values are the mean of three replicates; error is the standard error of the mean.

To determine whether Pmar-LigP and Ac42-Lig displayed similar fidelity trends with hachimodji DNA, we assayed singly mismatched substrates where AEGIS bases are positioned on the nick strand opposite canonical bases (Supplementary 10). Although these mismatch substrates sample only a small number of the 3248 possible combinations at the 5’ and 3’ nick termini, they are generated from recombining oligonucleotides used for fully-matched ligation assays making these experiments more directly comparable.

In contrast with the strong discrimination against canonical mismatches at the 3’ terminus, both Pmar-LigP and Ac42-Lig joined AEGIS mismatches with high efficiency for all tested combinations at either terminus, even under the standard (30 min at 25 °C, 20 nM enzyme, no additives) reaction conditions (Figure 9 Bi), and this lack of fidelity was further amplified under the optimised (18 h 15 °C, excess enzyme, 10% PEG) condition (Figure 9 Bii). To validate this result, we generated symmetrical mismatches where the AEGIS base is in the complement strand with an opposing natural mismatch in the ligatable nick (Supplementary 10). This showed a similar trend with all 3’ mismatches readily ligated under the optimised condition, and both the 3’ A:**pS** and 3’ A:**Z** joined under the standard reaction condition.

**Figure 9.**
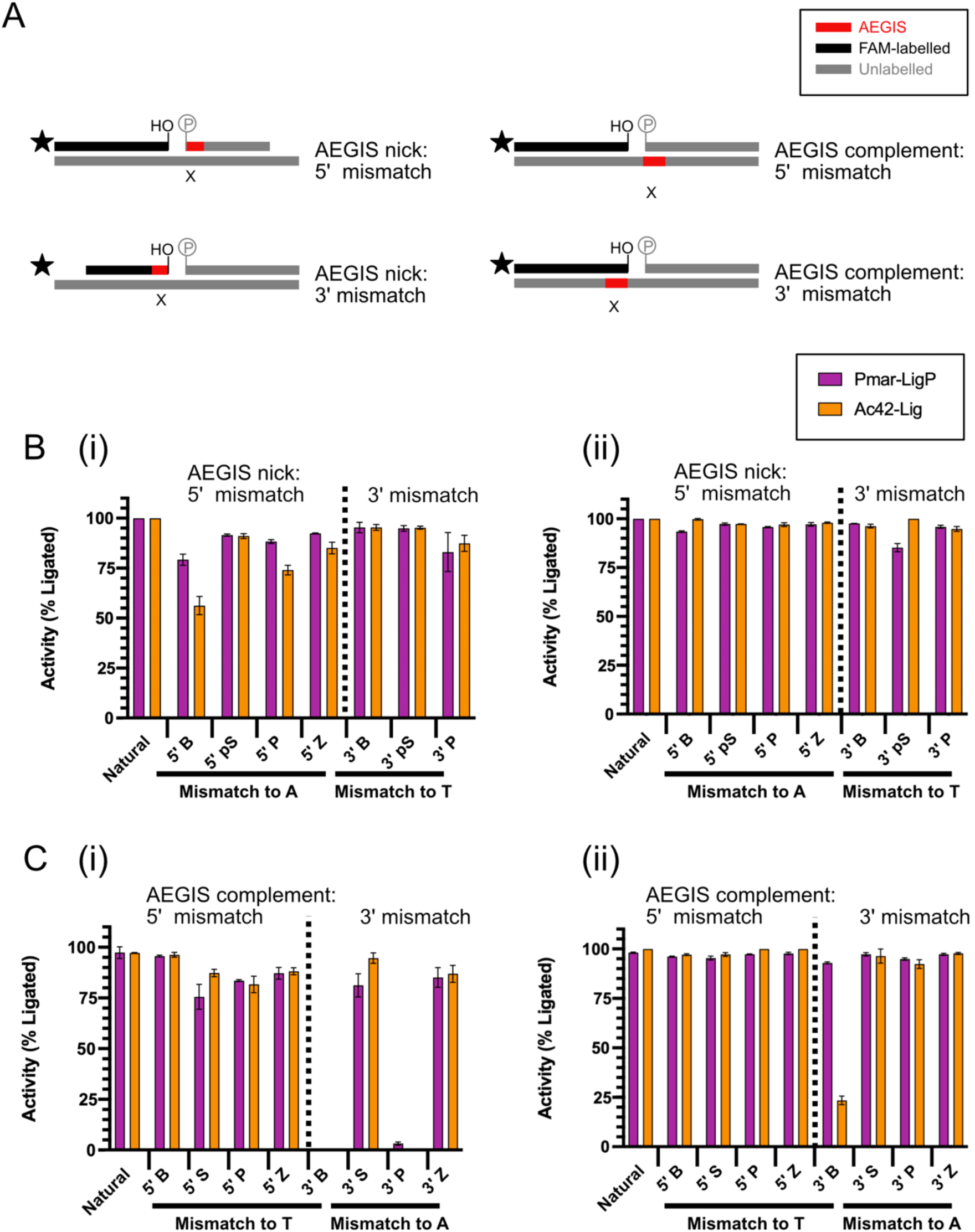
Ligation of single mismatched hachimoji bases by Pmar-LigP and Ac42-Lig. A) Schematic of DNA substrates containing single AEGIS-canonical base pair mismatches; oligonucleotides and duplex sequences are given in supplementary sections 2 and 10 respectively. B) ligation yield with single mismatched bases where the AEGIS base is in the ligatable nick strand under standard conditions (20 nM enzyme and incubation time 30 min at 25 °C) (i) and optimised conditions (10% PEG 3350, 1.9μM enzyme and incubation time 18 hours at 15 °C) (ii). C) ligation yield with single mismatched bases where the AEGIS base is in the complement strand under standard conditions (i) and optimised conditions (ii). All values are the mean of three replicates and error is the standard error of the mean.

Given this significant low-fidelity ligation of single AEGIS mismatches, we tested if substrates where both the 5’ and 3’ termini contained an AEGIS base could be joined when mis paired with canonical nucleobases. By combining the ligatable oligonucleotides from the ‘AEGIS both sides’ series with a fully canonical complement, and conversely the ‘AEGIS both sides’ complement strand with canonical nicked oligonucleotides (Supplementary 10), we generated a mismatched substrate series where the single AEGIS base at each terminus was mis paired against a canonical (3’T, 5’A) in the complement (Figure 10 A), or a nick with a 3’A and 5’T was mis paired against each AEGIS base (Figure 10 B). Even under standard conditions (30 min at 25 °C, 20 nM enzyme, no additives), both Pmar-LigP and Ac42-Lig showed some low-fidelity joining of the double-mismatches including a 3’ A:**Z** joined to all 5’ mismatched termini tested, and 3’ **P**:T being joined when juxtaposed with 5’ **Z**:A and **S**:A mismatches. Again, the optimised AEGIS ligation conditions (18 h 15 °C, excess enzyme, 10% PEG) further enhanced promiscuous ligation with Ac42-Lig joining all combinations to some extent, and Pmar-LigP retaining discrimination only against mismatches **pS** base at the 3’ end.

**Figure 10.**
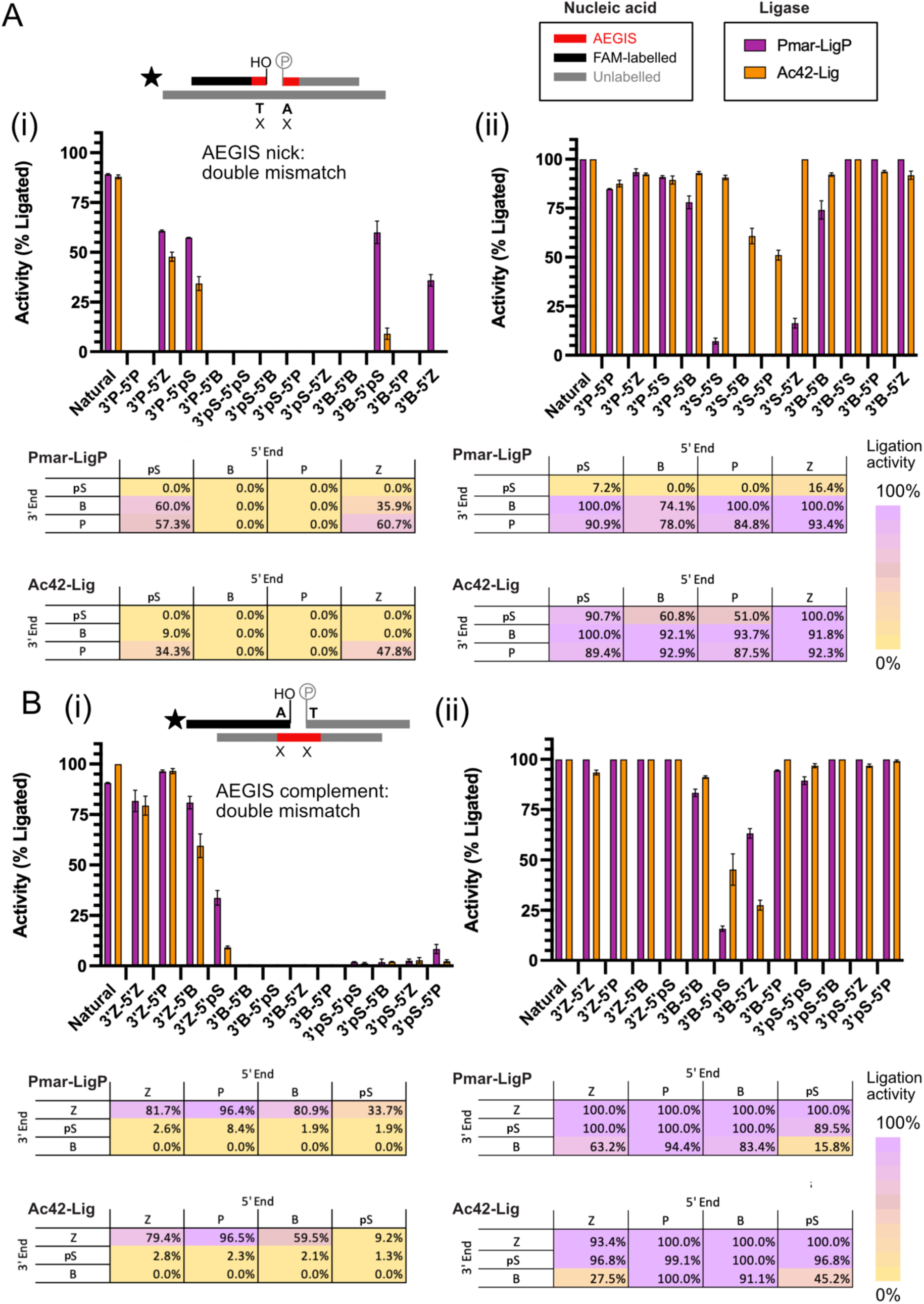
Ligation of mismatched hachimoji bases on both sides of the DNA nick by Pmar-LigP and Ac42-Lig. oligonucleotides and duplex sequences are given in supplementary sections 2 and 10 respectively. A) Ligation of substrates where the mismatched AEGIS base is in the nicked strand assayed under standard conditions (20 nM enzyme and incubation time 30 min at 25 °C) (i) or optimised conditions (10% PEG 3350, 1.9μM enzyme and incubation time 18 hours at 15 °C) (ii). B) Ligation of substrates where the mismatched AEGIS base is in the complement strand assayed under standard conditions (i) or optimised conditions (ii). All values are the mean of three replicates; error is the standard error of the mean.

To determine if this low-fidelity joining also occurs with consecutive AEGIS bases, we constructed substrates where these were mis paired against canonical bases on each terminus separately, or on both sides of the nick (Supplementary 10). Under standard assay conditions, only the 5’ nick **ZZZ** substrate was ligated by Ac42-Lig (less than 25%), while no other multiple AEGIS mismatches were joined to any observable extent (Figure 11 B(i) and C(i)). Under optimised reaction conditions however, Ac42-Lig joined all substrates with consecutive AEGIS-containing mismatches in the 5’ end of the nick, while Pmar-LigP effectively joined mismatches when the mismatched AEGIS bases are in the complement and to a lesser extent (less than 20%) when AEGIS bases were on the nicked strand (Figure 11 B(ii)). There is minimal ligation of consecutive 3’ AEGIS mismatches by either enzyme, but interestingly, both enzymes had high activity where six consecutive **Z** and **P** bases are positioned in the complement strand (Figure 11 C(ii)).

**Figure 11.**
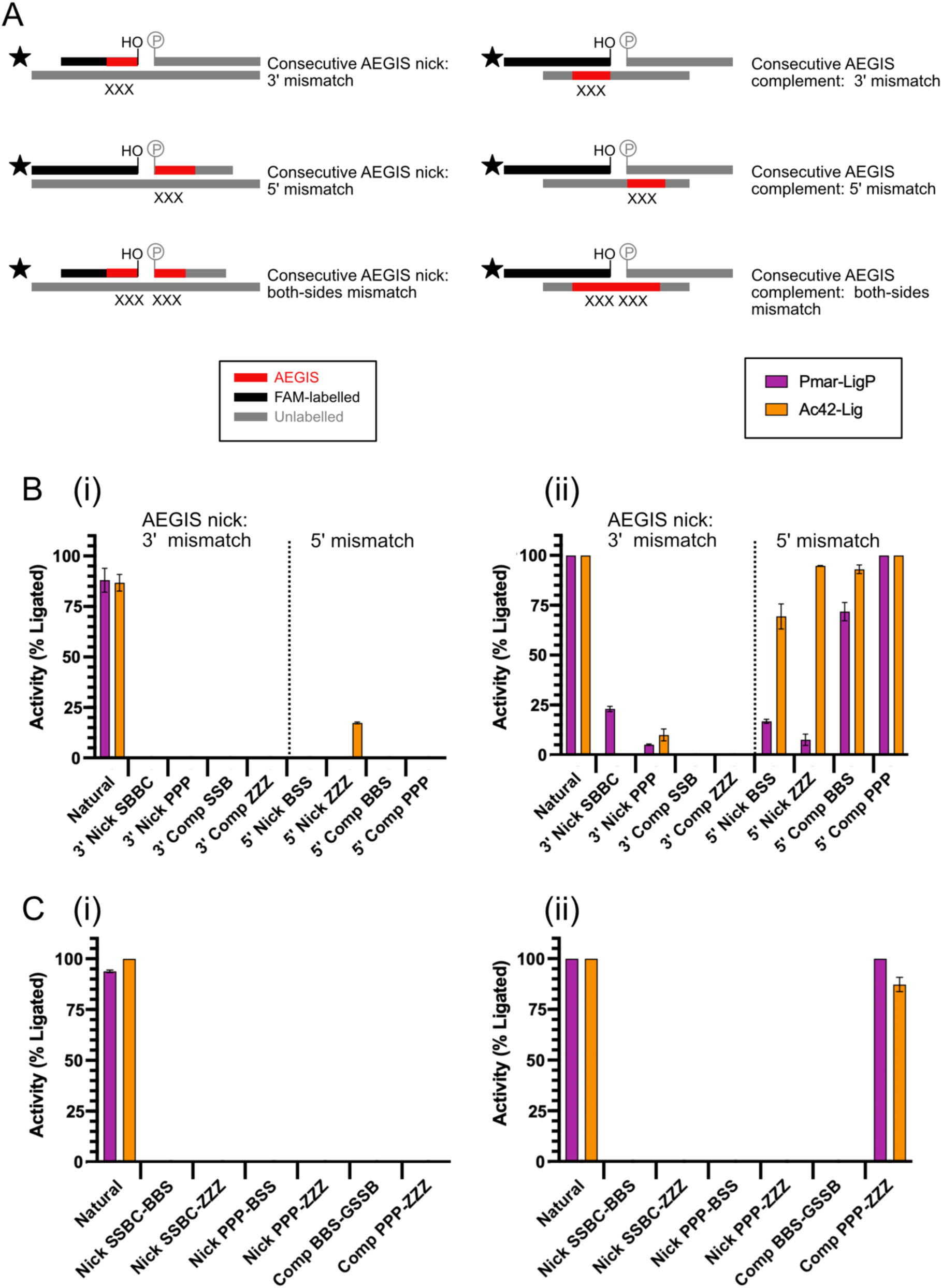
Ligation of consecutive mismatched AEGIS bases by Pmar-LigP and Ac42-Lig. A) Schematics of hachimoji substrates; oligonucleotides and duplex sequences are given in supplementary sections 2 and 10 respectively. B) Ligation of substrates where consecutive mismatched AEGIS are at either the 3’ or the 5’ terminus. Assays were run under standard conditions (20 nM enzyme and incubation time 30 min at 25 °C) (i) or optimised conditions (10% PEG 3350, 1.9μM enzyme and incubation time 18 hours at 15 °C) (ii). C) Ligation of substrates where consecutive mismatched AEGIS are at both termini. Assays were run under standard conditions (i) or optimised conditions. All values are the mean of three replicates; error is the standard error of the mean.

## DISCUSSION

Here we evaluate the activity and fidelity of DNA ligases when joining breaks in duplexes that were built from an eight-nucleotide hachimoji DNA alphabet. Our initial objective was to determine whether DNA ligases can join AGEIS-containing nicks to the same extent as natural DNA, and whether ligase structure (‘minimal’ or globular DB-domain) or AEGIS base composition (position, number and type) impacts ligation efficiency. We find the most significant determinants of hachimoji ligation are the number and positions of the non-standard base pairs, with all enzymes tested having a considerable bias against joining breaks with consecutive non-standard pairs at the 3’ acceptor end. Further, under standard reaction conditions (30 min at 25 °C, 20 nM enzyme, no additives) only Pmar-LigP and Ac42-Lig, both of which have globular DB-domains, exhibit any ligation activity with consecutive non-standard pairs on both sides of the break.

The decreased ligation yield when AEGIS base pairs are positioned at the 3’ side of the break is consistent with biases in ligation with 3’ hydrophobic pairs (7) as well as higher discrimination against mismatches at this position (7,25,26). However, the magnitude of this effect with hachimoji DNA, and especially the impact of having multiple AEGIS bases spanning the break, was unexpected. DNA ligases have not evolved to interrogate the duplex sequence surrounding a DNA break and, provided adjacent bases are correctly matched, most show minimal sequence bias which is consistent with their biological roles in joining replication and repair intermediates in a range of sequence contexts (6,26). Given **S:B** and **P:Z** pairs present helical parameters similar to those of A:T and G:C pairs and have a canonical DNA sugar-phosphate backbone, we had anticipated that even consecutive tracts of these non-canonical hachimoji bases should be well-tolerated by natural DNA ligases.

Numerous crystal structures of ligase enzymes in complex with nicked (natural) DNA, including most of the enzymes used in this study (Supplementary figure 11), have shown there are no specific contacts between the nucleotide bases and the enzyme, and the majority of the interactions are with the non-bridging phosphate oxygens (11,12,16–19). Key structural differences between the AEGIS nucleobases and their natural counterparts which may explain this discrimination include substitution of the 5-position of **S** and **pS**, (methyl) and **Z** (nitro) which would be situated in the major groove of the duplex. Meanwhile, although the **Z:P** pair presents similar minor groove functionalities to the C:G pair with a keto-group on the smaller (pyrimidine-analogue) ring and an amino group on the larger (purine-analogue) ring, this configuration is reversed in the **S:B** pair. These minor groove differences in particular may account for decreased ligation as previous work using modified nucleobases indicates that minor groove modification causes defective ligase binding (6). Analysis of hachimoji substrate binding by Pmar-LigP and Ac42-Lig, however, shows that for several key substrates, including DNA with consecutive AEGIS bases at the termini, substrate affinity is not significantly decreased. This suggests that for the consecutive AEGIS-terminated nicks examined here, catalytic steps rather than substrate engagement are impacted. DNA engagement (natural or AEGIS) requires the DNA ligase to be covalently adenylated (step 1 of the ligation reaction). In our experiments recombinant ligase was preincubated with ATP during purification meaning the adenylation status of the ligase would be equivalent for all samples. As enzyme re-adenylation is independent of the nucleic acid substrate, step 1 differences would not explain any discrepancies in hachimoji DNA binding or activity. Instead, step 2 (adenyl transfer from the enzyme to the DNA), step 3 (nucleophilic attack of the adenylated 5’ terminus on the 3’ OH) or both are likely affected. Previous work using long timescale molecular dynamics simulations indicates that hachimoji DNA containing consecutive **P:Z** bases has altered solution dynamics which arise from differences in the electrostatic properties of these nucleobases resulting in widening of the major groove of the duplex (14). With DNA polymerases, these altered dynamics of AEGIS DNA explain the bias of wild-type Klentaq against templated-direct incorporation of **Z** nucleotides, and may have a similar impact of DNA ligase activity (27). Although previous studies of hachimoji DNA duplexes crystalised with a host protein (Moloney murine leukemia virus reverse transcriptase) show the helical conformation of DNA is minimally affected by inclusion of AEGIS bases (3), this may not apply at the less-constrained ends of the substrate nick. Similarly, it is possible that sequential AEGIS bases interfere with the ligase-induced distortion of the duplex at the nick site which is seen in all crystal structures of ligase bound to natural DNA (11,12,16–19), and is necessary for ligation to occur.

In other hachimoji sequence contexts with single AEGIS termini, disrupted binding did appear to be responsible for low activity, especially for Ac42-Lig where the 3’ terminal nucleobase of the nick was a **B** and the 5’ terminus was another AEGIS base. One limitation of the current study is all single-AEGIS terminal combinations were tested with the same canonical flanking sequences, and it would be useful to investigate whether strong biases against some AEGIS combinations remain in a wider range of canonical sequence contexts, especially where binding is markedly affected. Previous papers have noted a “complex relationship between ligase identity and fidelity” when investigating mismatch ligation (26), and we observe a similar phenomenon in the outcomes ligation with AEGIS termini reported here . Despite the ligases we tested showing broad trends such as discrimination against consecutive AEGIS bases at the 3’ terminus, the details of AEGIS ligation vary with individual ligase. Pmar-LigP and Ac42-Lig both have broadly similar structures, but show distinct biases against certain AEGIS bases, especially in the 3’ position.

Previous work where DNA ligase was used to incorporate AEGIS bases into hachimoji DNA used conditions similar to the optimised reaction conditions reported here to effect joining of termini containing multiple AEGIS bases (5). Addition of molecular crowding agents such as PEG is a common strategy to enhance ligation of other challenging substrates including ‘blunt’ (non-cohesive double-stranded breaks) single base-pair overhangs (e.g. T-A cloning) as well as nucleic acids with modified backbones (28–30). However, this increase in joining efficiency must be balanced against a potential decrease in fidelity as previous studies show that crowding agents alongside high ligase concentrations and extended incubation times promote ligation of mismatched bases pairs with natural DNA (8). Here we find that even without the addition of crowding agents, AEGIS mismatches are joined with high frequency, and this fidelity is even lower when PEG is added in combination with high enzyme concentrations and long incubation times. The notably low-fidelity ligation of AEGIS 3’ mismatches, which contrasts with the stringency observed in the purely canonical set, may arise from potential of some AEGIS bases to mis pair with canonical counterparts due to tautomerisation or deprotonation (31). Here, the enol tautomer of **B** mis pairs with T and the keto imino tautomer of **B** mis pairs with A, which would account for the ready ligation of these mismatches at either terminus. Similarly, a minor tautomer of **Z** can form two hydrogen bonds with A closely resembling a native A:T pair (supplementary 12). Although the T:**P** mismatch has only two of three possible hydrogen bonds, its pyrimidine:purine pairing could be accommodated in the helix geometry and this mismatch has been observed during PCR incorporation by Taq polymerase (32). It is also possible that the environment of the ligase active site stabilises tautomeric forms of AEGIS bases promoting low-fidelity outcomes. Recent structural snapshots have defined the molecular basis for mismatch discrimination by human ligase I (33), and structural studies of AEGIS-canonical mismatches in complex with DNA ligase would provide similarly useful information on permissibility of joining. The two 3’ combinations which were not joined under the standard condition (30 min at 25 °C, 20 nM enzyme, no additives), A:**B** and A:**P** have both steric (bulky purine:purine pair) and electrochemical features which make pairing and hence ligation highly unfavourable. All AEGIS termini which are ligated in a mismatch context were also ligated with high efficiency in the correctly matched substrates indicating that mismatch to canonical nucleobases was not promoting ligation of AEGIS termini which would not have otherwise being joined.

Despite only investigating a sub-set of the possible AEGIS-canonical mismatch combinations, this finding of low-fidelity joining highlights a potential challenge in using ligation-based strategies for AEGIS base incorporation into DNA duplexes. Investigation of wider sequence contexts and mismatch combinations will be important for future synthetic biology endeavours as well as understanding mechanistic aspects of accurate and mismatched ligation. Similarly, determining DNA ligase efficiency and fidelity with AEGIS-containing cohesive-ended substrates will be informative for synthetic workflows. To practically interrogate the enormous number of sequence combinations, high throughput strategies based on capillary electrophoresis (7) or sequencing (34,35) will be necessary, with the latter approach being made more feasible by recent advances in AEGIS base sequence determination (5,32).

The recent success of several ligation-dependent methods for synthesising modified nucleic acids has underscored the utility of these enzymes to deliver products which are either unachievable or inefficient using solid-phase or polymerase-mediated strategies (5,36–38). In a recent proof-of-concept, Kawabe *et al.* used an ingenious three-enzyme strategy including a penultimate ligation step to install individual non canonical base pairs into DNA duplexes. This work showed incorporation of all four hachimoji pairs (**S:B**, **P:Z**, A:T and C:G) as well as two additional orthogonal hydrogen bonded base pairs **V:J** and **K:X** into duplex DNA; an outcome not accessible by chemical means due to the instability of precursors for phosphoramidite synthesis (5). Meanwhile, Sabat *et al*. have developed a flexible chemoenzymatic synthesis method for base and backbone-modified nucleic acid duplexes which uses a DNA complement strand as a template for programmable recruitment of ‘shortmer’ base-modified oligonucleotides which are then assembled into the final polymer by enzymatic ligation (36). Similarly, T4 DNA ligase has been used to ligate duplexes containing single hydrophobic base pairs (39) as well as being deployed to join a wide range of non-canonical nucleic acids with modified sugars, representing a pathway for production of mixed-chemistry backbones (30,40–42).

To date, DNA ligases deployed in synthesis of modified nucleic acids, including hachimoji DNA, have been wild-type commercially available enzymes; predominantly T3 and T4 DNA ligases. Probing the intrinsic capacity of a broad range of wild-type ligases to join non-canonical nucleobase DNA has the potential to not only deliver new enzymes for biotechnological purposes, but also to improve our understanding of substrate recognition and end-discrimination by natural DNA ligase enzymes. For synthetic biology, engineered ligases capable of joining AEGIS bases with better efficiency and fidelity would be desirable, and such variants would provide fundamental insight into the evolutionary landscape of DNA ligases toward non-natural substrates.

