## Supplement for "DNA Ligases Discriminate Between Natural and Non-Natural Base Pairs"

#### Supplementary 1

Summary of DNA ligases used in this study

| Ligase name | Originating organism | Reference |
|---|---|---|
| Bsp-Lig | *Burkholderia pseudomallei* | (Pan, Lian et al. 2021) |
| Pmar-LigP | *Prochlorococcus marinus* | (Williamson and Leiros 2019) |
| Ame-Lig | *Alteromonas mediterranea* | (Williamson, Grgic et al. 2018) |
| Pmar-LigW | *Prochlorococcus marinus* | (Williamson, Hjerde et al. 2016) |
| Ac42-Lig | *Acinetobacter* phage Ac42 | (Rothweiler, Leiros et al. 2025) |
| P10VF-Lig | *Rhizobium* phage vB_RleM_P10VF | (Rothweiler, Leiros et al. 2025) |

#### Supplementary 2

A) Oligonucleotide sequences used to generate substrates

| **Oligonucleotide name** | **Sequence (5' – 3')** | **Modifications** |
|---|---|---|
| NL1 | AGGCCATGGCTGATATCGCA | 5' 6-fluorescein |
| NL2 | TAGGCATTCGAGCTCCGTCG | 5' phosphate |
| NL3 | CGACGGAGCTCGAATGCCTATGCGATATCAGCCATGGCCT | |
| NL8 | ATATCAGCCATGGCCT | 5' phosphate |
| NL9 | CGACGGAGCTCGAATGCCTATGCG | |
| NL10 | CGACGGAGCTCGAATGCCTACGCGATATCAGCCATGGCCT | |
| NL14 | CGACGGAGCTCGAATGCCTAGGCGATATCAGCCATGGCCT | |
| NL15 | CGACGGAGCTCGAATGCCTATGCGATATCAGCCATGGCCT | |
| NL16 | CGACGGAGCTCGAATGCCTATGCGATATCAGCCATGGCCT | |
| UB14 | TGGCTGATATC**PPP** | 5' 6-fluorescein |
| UB15 | AGCTCGAATGCCTA**ZZZ**GATATCAGCCA | |
| UB16 | **ZZZ**GCATTCGAGCT | 5' phosphate |
| UB17 | AGCTCGAATGC**PPP**TGCGATATCAGCCA | |
| UB18 | AGCTCGAATGC**PPPZZZ**GATATCAGCCA | |
| SB2 | **B**AGGCATTCGAGCT | 5' phosphate |
| SB14 | TGGCTGATAT**SBB**C | 5' 6-fluorescein |
| SB15 | AGCTCGAATGCCTAG**SSB**ATATCAGCCA | |
| SB16 | **BSS**GCATTCGAGCT | 5' phosphate |
| SB17 | AGCTCGAATGC**BBS**TGCGATATCAGCCA | |
| SB18 | AGCTCGAATGC**BBS**GSSBATATCAGCCA | |
| pSB14 | TGGCTGATAT**SBB** | 5' 6-fluorescein; S= pseudoC |
| pSB15 | AGCTCGAATGCCTAG**SSB**ATATCAGCCA | |
| pSB16 | **BSS**GCATTCGAGCT | 5' phosphate; S= pseudoC |
| pSB17 | AGCTCGAATGC**BBS**TGCGATATCAGCCA | S= pseudoC |
| pSB18 | AGCTCGAATGC**BBSSSB**ATATCAGCCA | S= pseudoC |
| NL2-v2 | CTAGGCATTCGAGCTCCGTCG | 5' phosphate |
| UB-Ni1 | TGGCTGATATCGC**P** | 5' 6-fluorescein |
| UB-Ni2 | **P**AGGCATTCGAGCT | 5' Phosphate |
| UB-Ni3 | AGCTCGAATGCCT**ZZ**GCGATATCAGCCA | |
| UB-Ni4 | **Z**AGGCATTCGAGCT | 5' Phosphate |
| UB-Ni5 | AGCTCGAATGCCT**PZ**GCGATATCAGCCA | |
| UB-Ni6 | **S**AGGCATTCGAGCT | 5' Phosphate; S= pseudoC |
| UB-Ni7 | AGCTCGAATGCCT**BZ**GCGATATCAGCCA | |
| UB-Ni8 | AGCTCGAATGCCT**SZ**GCGATATCAGCCA | S= pseudoC |
| UB-Ni9 | TGGCTGATATCGC**S** | 5' 6-fluorescein; S= pseudoC |
| UB-Ni10 | AGCTCGAATGCCT**BB**GCGATATCAGCCA | |
| UB-Ni11 | AGCTCGAATGCCT**SB**GCGATATCAGCCA | S= pseudoC |
| UB-Ni12 | AGCTCGAATGCCT**ZB**GCGATATCAGCCA | |
| UB-Ni13 | AGCTCGAATGCCT**PB**GCGATATCAGCCA | |
| UB-Ni14 | TGGCTGATATCGC**B** | 5' 6-fluorescein |
| UB-Ni15 | AGCTCGAATGCCT**SS**GCGATATCAGCCA | S= pseudoC |
| UB-Ni16 | AGCTCGAATGCCT**BS**GCGATATCAGCCA | S= pseudoC |
| UB-Ni17 | AGCTCGAATGCCT**ZS**GCGATATCAGCCA | S= pseudoC |
| UB-Ni18 | AGCTCGAATGCCT**PZ**SGCGATATCAGCCA | S= pseudoC |
| UB-OH1 | TGGCTGATATPP**ZZZP** | 5' 6-fluorescein |

| | | |
|---|---|---|
| UB-OH2 | **Z**PCATTCGAGCT | 5’ Phosphate |
| UB-OH3 | **ZZ**ATATCAGCCA | 5’ Phosphate |
| UB-OH4 | AGCTCGAATG**ZPZPPP** | |
| UB-OH5 | TGGCTGATA**PPP**CGCA | 5' 6-fluorescein |
| UB-OH6 | **ZZZ**GATATCAGCCA | 5’ Phosphate |
| UB-OH7 | CTCGAATGC**BBS**TGCG | S= pseudoC |
| MM1 | AGGCCATGGCTGATATCATC | 5' 6-fluorescein |
| MM2 | CGAGCATTCGAGCTCCGTCG | 5’ phosphate |
| MM3 | CGACGGAGCTCGAATGCCT**B**TGCGATATCAGCCATGGCCT | |
| MM4 | CGACGGAGCTCGAATGCCT**S**TGCGATATCAGCCATGGCCT | |
| MM5 | CGACGGAGCTCGAATGCCT**P**TGCGATATCAGCCATGGCCT | |
| MM6 | CGACGGAGCTCGAATGCCT**Z**TGCGATATCAGCCATGGCCT | |
| MM7 | CGACGGAGCTCGAATGCCTA**B**GCGATATCAGCCATGGCCT | |
| MM8 | CGACGGAGCTCGAATGCCTA**S**GCGATATCAGCCATGGCCT | |
| MM9 | CGACGGAGCTCGAATGCCTA**P**GCGATATCAGCCATGGCCT | |
| MM10 | CGACGGAGCTCGAATGCCTA**Z**GCGATATCAGCCATGGCCT | |

B) Oligonucleotide combinations used to generate substrates

| **Substrate name** | **Acceptor (FAM; 3’ OH of nick)** | **Donor (5’ phosphate of nick)** | **Complement** |
|---|---|---|---|
| **Control** | | | |
| Nicked | NL1 | NL2 | NL3 |
| **Consecutive AEGIS bases** | | | |
| 3’ Acceptor P:Z | UB14 | NL2 | UB15 |
| 5’ Donor P:Z | NL1 | UB16 | UB17 |
| Both Sides P:Z | UB14 | UB16 | UB18 |
| 3’ Acceptor S:B | SB14 | NL2 | SB15 |
| 5’ Donor S:B | NL1 | SB16 | SB17 |
| Both Sides S:B | SB14 | SB16 | SB18 |
| 3’ Acceptor pS:B | pSB14 | NL2-v2 | pSB15 |
| 5’ Donor pS:B | NL1 | pSB16 | pSB17 |
| Both Sides pS:B | pSB14 | pSB16 | pSB18 |
| **Single AEGIS bases** | | | |
| 3’P-5’P | UB-Ni1 | UB-Ni2 | UB-Ni3 |
| 3’P-5’Z | UB-Ni1 | UB-Ni4 | UB-Ni5 |
| 3’P-5’S | UB-Ni1 | UB-Ni6 | UB-Ni7 |
| 3’P-5’B | UB-Ni1 | SB2 | UB-Ni8 |
| 3’S-5’S | UB-Ni9 | UB-Ni6 | UB-Ni10 |
| 3’S-5’B | UB-Ni9 | SB2 | UB-Ni11 |
| 3’S-5’P | UB-Ni9 | UB-Ni2 | UB-Ni12 |
| 3’S-5’Z | UB-Ni9 | UB-Ni4 | UB-Ni13 |
| 3’B-5’B | UB-Ni14 | SB2 | UB-Ni15 |
| 3’B-5’S | UB-Ni14 | UB-Ni6 | UB-Ni16 |
| 3’B-5’P | UB-Ni14 | UB-Ni2 | UB-Ni17 |
| 3’B-5’Z | UB-Ni14 | UB-Ni4 | UB-Ni18 |
| **Natural Mismatches** | | | |
| 3’ A:C | NL1 | NL2 | NL10 |
| 3’ A:G | NL1 | NL2 | NL14 |
| 5’ C:A | NL1 | NL2 | NL15 |
| 5’ G:A | NL1 | NL2 | NL16 |
| **AEGIS nick- single mismatch** | | | |
| 5’ B:A Nick Mismatch | NL1 | SB2 | NL3 |
| 5’ S:A Nick Mismatch | NL1 | UB-Ni6 | NL3 |
| 5’ P:A Nick Mismatch | NL1 | UB-Ni2 | NL3 |

| | | | |
|---|---|---|---|
| 5’ Z:A Nick Mismatch | NL1 | UB-Ni4 | NL3 |
| 3’ B:T Nick Mismatch | UB-Ni14 | NL2 | NL3 |
| 3’ S:T Nick Mismatch | UB-Ni9 | NL2 | NL3 |
| 3’ P:T Nick Mismatch | UB-Ni1 | NL2 | NL3 |
| **AEGIS complement- single mismatch** | | | |
| 5’ T:B Comp Mismatch | NL1 | NL2 | MM3 |
| 5’ T:S Comp Mismatch | NL1 | NL2 | MM4 |
| 5’ T:P Comp Mismatch | NL1 | NL2 | MM5 |
| 5’ T:Z Comp Mismatch | NL1 | NL2 | MM6 |
| 3’ A:B Comp Mismatch | NL1 | NL2 | MM7 |
| 3’ A:S Comp Mismatch | NL1 | NL2 | MM8 |
| 3’ A:P Comp Mismatch | NL1 | NL2 | MM9 |
| 3’ A:Z Comp Mismatch | NL1 | NL2 | MM10 |
| **AEGIS nick- double mismatch** | | | |
| P-P Nick Mismatch | UB-Ni1 | UB-Ni2 | NL3 |
| P-Z Nick Mismatch | UB-Ni1 | UB-Ni4 | NL3 |
| P-S Nick Mismatch | UB-Ni1 | UB-Ni6 | NL3 |
| P-B Nick Mismatch | UB-Ni1 | SB2 | NL3 |
| S-S Nick Mismatch | UB-Ni9 | UB-Ni6 | NL3 |
| S-B Nick Mismatch | UB-Ni9 | SB2 | NL3 |
| S-P Nick Mismatch | UB-Ni9 | UB-Ni2 | NL3 |
| S-Z Nick Mismatch | UB-Ni9 | UB-Ni4 | NL3 |
| B-B Nick Mismatch | UB-Ni14 | SB2 | NL3 |
| B-S Nick Mismatch | UB-Ni14 | UB-Ni6 | NL3 |
| B-P Nick Mismatch | UB-Ni14 | UB-Ni2 | NL3 |
| B-Z Nick Mismatch | UB-Ni14 | UB-Ni4 | NL3 |
| **AEGIS complement- double mismatch** | | | |
| Z-Z Comp Mismatch | NL1 | NL2 | UB-Ni3 |
| Z-P Comp Mismatch | NL1 | NL2 | UB-Ni5 |
| Z-B Comp Mismatch | NL1 | NL2 | UB-Ni7 |
| Z-S Comp Mismatch | NL1 | NL2 | UB-Ni8 |
| B-B Comp Mismatch | NL1 | NL2 | UB-Ni10 |
| B-S Comp Mismatch | NL1 | NL2 | UB-Ni11 |
| B-Z Comp Mismatch | NL1 | NL2 | UB-Ni12 |
| B-P Comp Mismatch | NL1 | NL2 | UB-Ni13 |
| S-S Comp Mismatch | NL1 | NL2 | UB-Ni15 |
| S-B Comp Mismatch | NL1 | NL2 | UB-Ni16 |
| S-Z Comp Mismatch | NL1 | NL2 | UB-Ni17 |
| S-P Comp Mismatch | NL1 | NL2 | UB-Ni18 |
| **Consecutive AEGIS separate termini** | | | |
| 3’ nick SBBC mismatch | SB14 | NL2 | NL3 |
| 5’ nick BSS mismatch | NL1 | SB16 | NL3 |
| 3’ nick PPP mismatch | UB14 | NL2 | NL3 |
| 5’ nick ZZZ mismatch | NL1 | UB16 | NL3 |
| 3’ complement GSSB mismatch | NL1 | NL2 | SB15 |
| 5’ complement BBS mismatch | NL1 | NL2 | SB17 |
| 3’ complement ZZZ mismatch | NL1 | NL2 | UB15 |
| 5’ complement PPP mismatch | NL1 | NL2 | UB17 |
| **Consecutive AEGIS both termini** | | | |
| Nick SSBC-BBS mismatch | SB14 | SB16 | NL3 |
| Nick SSBC-ZZZ mismatch | SB14 | UB16 | NL3 |
| Nick PPP-BSS mismatch | UB14 | SB16 | NL3 |
| Nick PPP-ZZZ mismatch | UB14 | UB16 | NL3 |
| Complement BBS-GSSB mismatch | NL1 | NL2 | SB18 |
| Complement PPP-ZZZ mismatch | NL1 | NL2 | UB18 |

### Supplementary 3

SV-AUC *c(s)* distributions of FAM labelled natural/hachimoji substrates (0.82–1.0 uM) incubated with a concentration range of Pmar-LigP (0.18–6.2 uM). All plots were generated in GUSSI V 2.1.0. A is natural DNA; B is S:B (3’ acceptor); C is S:B (5’ donor); and D is S:B (both sides). Left pane: Continuous *c(s)* distributions were integrated using the GUSSI integration tool: shaded in blue is the natural/hachimoji DNA and shaded in red is the Pmar-LigP bound with the natural/hachimoji DNA. Middle pane: One representative fit of the *c(s)* model to raw data is shown for each natural/hachimoji DNA experiment. Right pane: Overall, all fits to the data were good, as indicated by the r.m.s.d. plots (Table). The r.m.s.d. relating to the fit in the middle pane is highlighted in red.

A

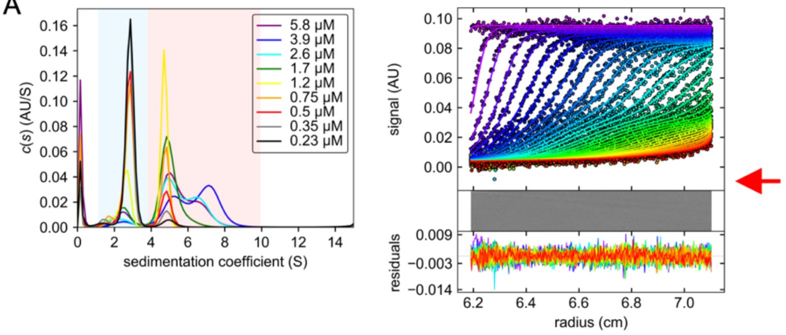


| [LigP] μM | RMSD |
|---|---|
| 5.8 | 0.0022 |
| 3.9 | 0.0022 |
| 2.6 | 0.0022 |
| 1.7 | 0.0020 |
| **1.2** | **0.0021** |
| 0.75 | 0.0022 |
| 0.5 | 0.0022 |
| 0.35 | 0.0022 |
| 0.23 | 0.0020 |

B

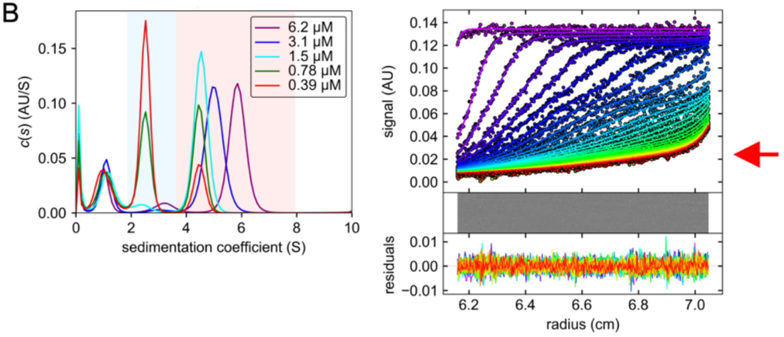


| [LigP] μM | r.m.s.d. |
|---|---|
| 6.2 | 0.0024 |
| 3.1 | 0.0023 |
| 1.5 | 0.0024 |
| **0.78** | **0.0021** |
| 0.39 | 0.0020 |

C

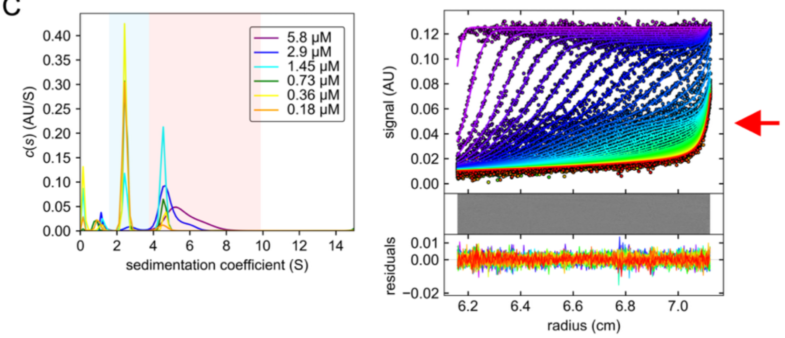


| [LigP] μM | r.m.s.d. |
|---|---|
| 5.8 | 0.0023 |
| 2.9 | 0.0024 |
| **1.45** | **0.0024** |
| 0.73 | 0.0024 |
| 0.36 | 0.0022 |
| 0.18 | 0.0022 |

D

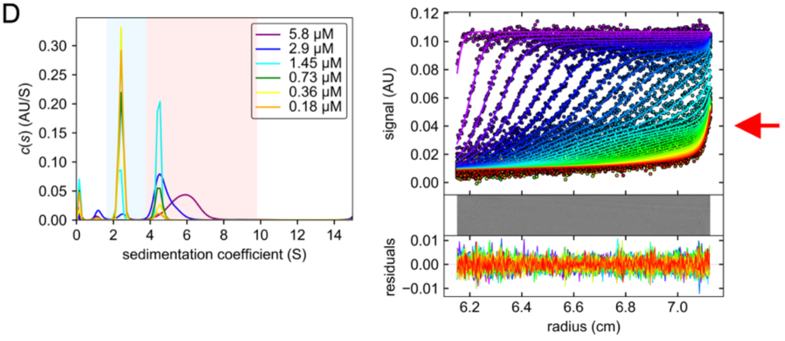


| [LigP] μM | r.m.s.d. |
|---|---|
| 5.8 | 0.0031 |
| 2.9 | 0.0023 |
| **1.45** | **0.0024** |
| 0.73 | 0.0024 |
| 0.36 | 0.0022 |
| 0.18 | 0.0021 |

Supplementary 4

DNA Ligase binding of Pmar-LigP and Ac42-Lig with hachimoji substrates evaluated by EMSA.

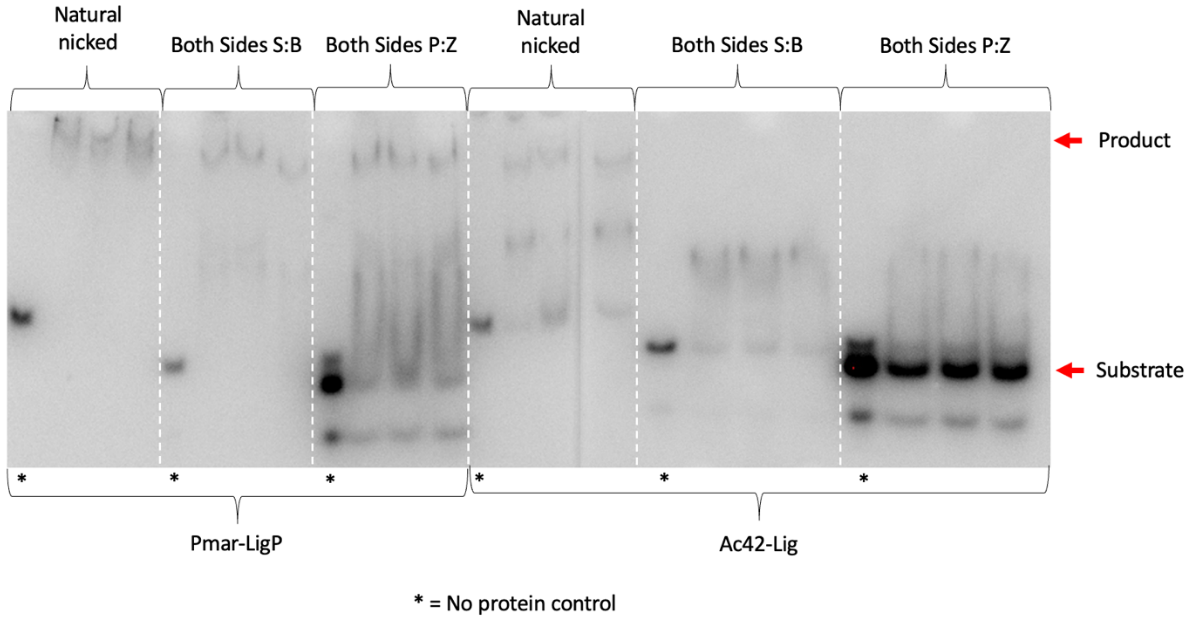


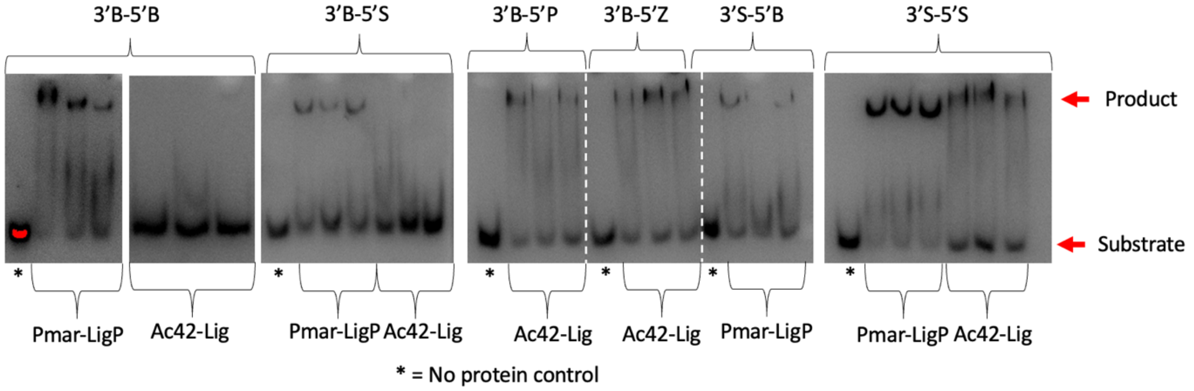

### Supplementary 5

Steady state kinetics of Pmar-LigP and Ac42-Lig with natural nicked DNA (A), 3'acceptor **S:B** (consecutive) hachimoji DNA (B) and 3'acceptor **P:Z** (consecutive) hachimoji DNA (C), and for Ac42-Lig with natural nicked DNA (D), 3'acceptor **S:B** (consecutive) hachimoji DNA (E) and 3'acceptor **P:Z** (consecutive) hachimoji DNA (F). Rates for each substrate concentration are plotted in panels (i), concentration of ligated product determined from urea-PAGE gels are shown in panel (ii). Enzyme concentrations are 10 nM for natural DNA, 20 nM for S:B and 20 nM for P:Z (Pmar-LigP) and 1nM natural, 20 nM for S:B and 20 nM for P:Z (Ac42-Lig). Urea PAGE gel images are available at DOI:10.5281/zenodo.20821393.

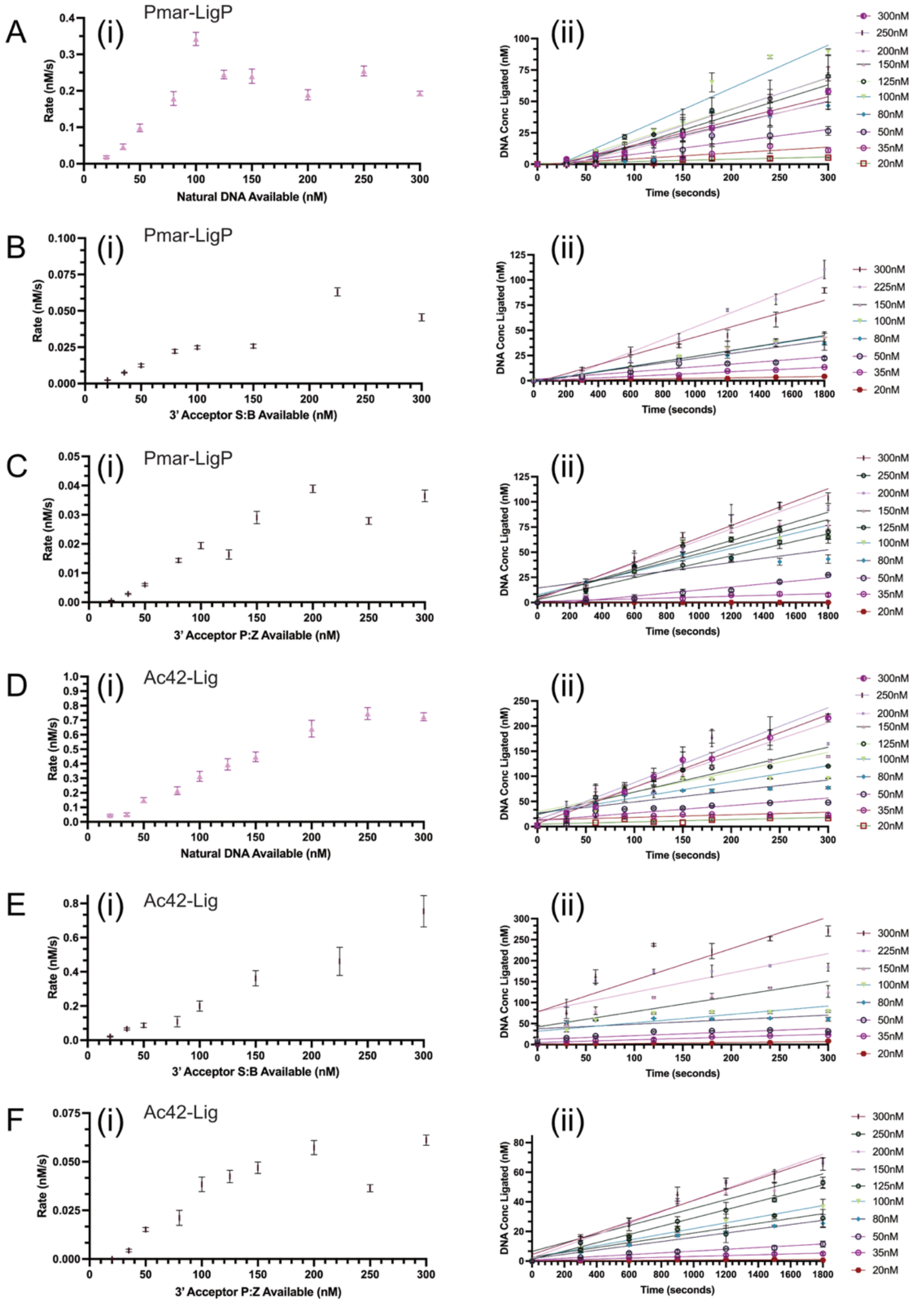

Supplementary 6

DNA Ligase activity of Pmar-LigP and Ac42-Lig with hachimoji substrates and varying ATP concentrations. Assays were run with 80nM double stranded DNA oligonucleotides, 10 mM $Mg^{2+}$, 50 mM TRIS pH 8.0, 50 mM NaCl, 1 mM DTT and 20nM of protein. ATP concentration varies between 0.1 mM to 5 mM. Samples are incubated at 25 °C for 30 minutes. Error bars are not visible for data points with standard deviation less than 3.0 %.

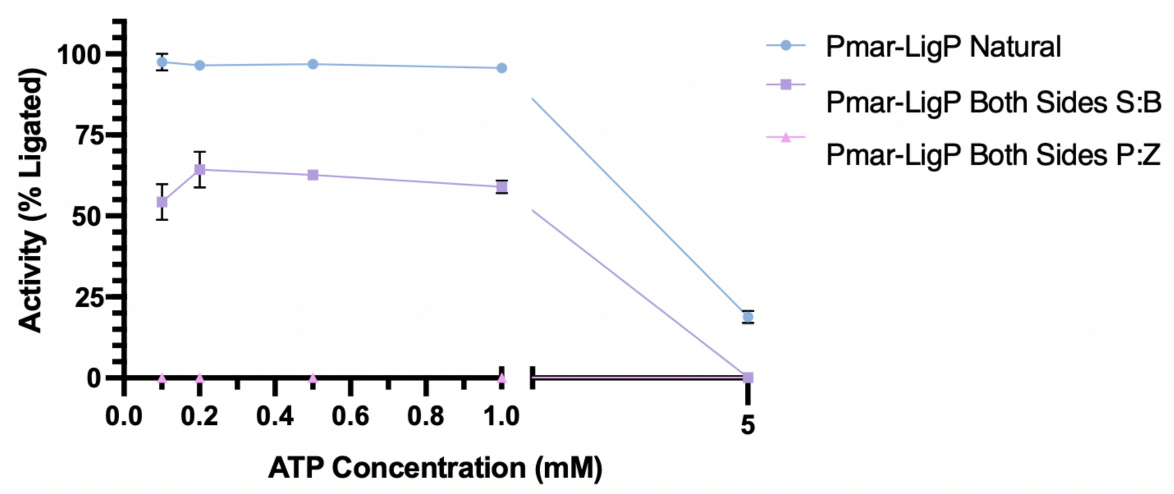


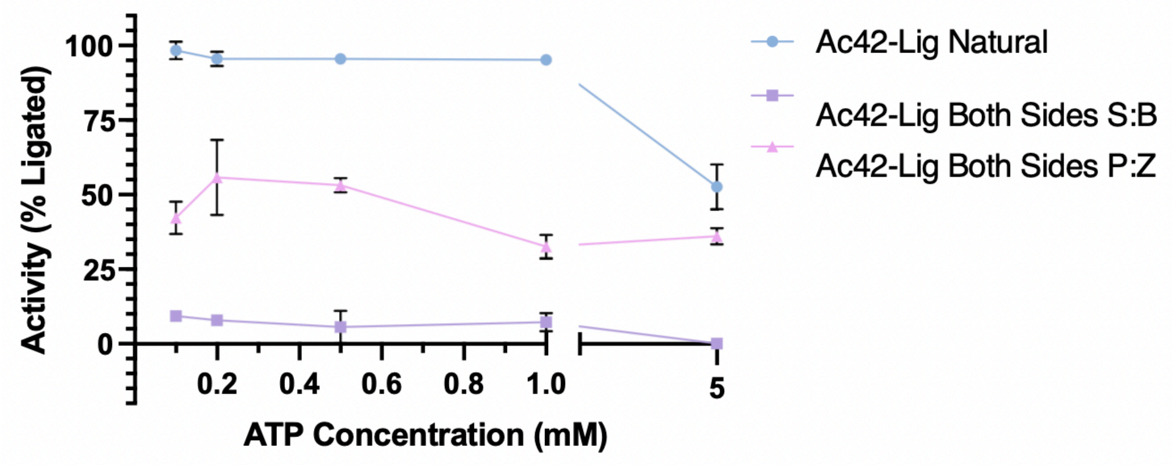

Supplementary 7

DNA Ligase activity of Pmar-LigP and Ac42-Lig with hachimoji substrates and varying magnesium ion concentration. Assays were run with 80 nM double stranded DNA oligonucleotides, 1 mM ATP, 50mM TRIS pH 8.0, 50 mM NaCl, 1 mM DTT and 20 mM of protein. $Mg^{2+,}$ concentration varies between 0.1 mM to 5 mM. Samples are incubated at 25 °C for 30 minutes. Error bars are not visible for data points with standard deviation less than 3.0 %.

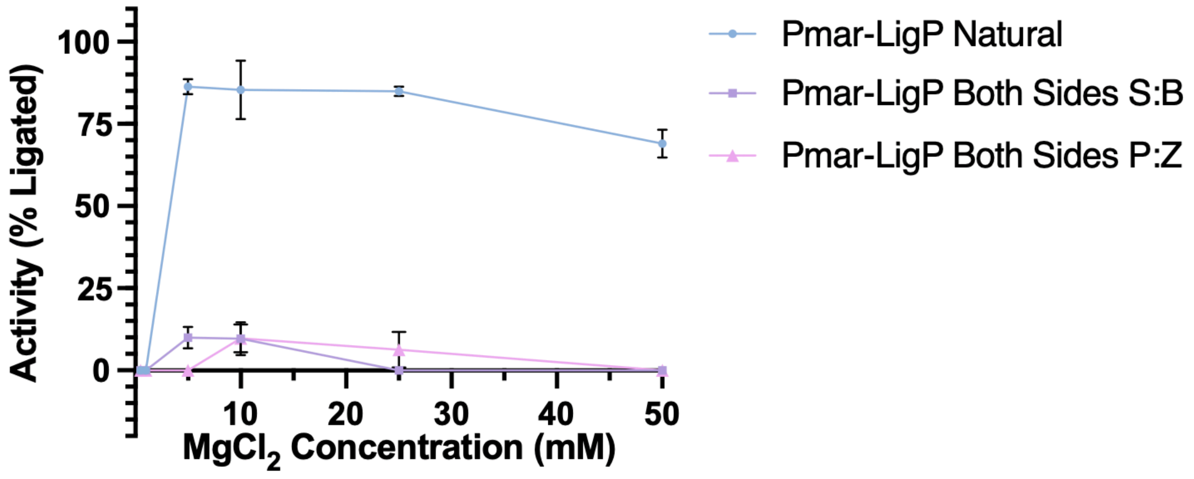


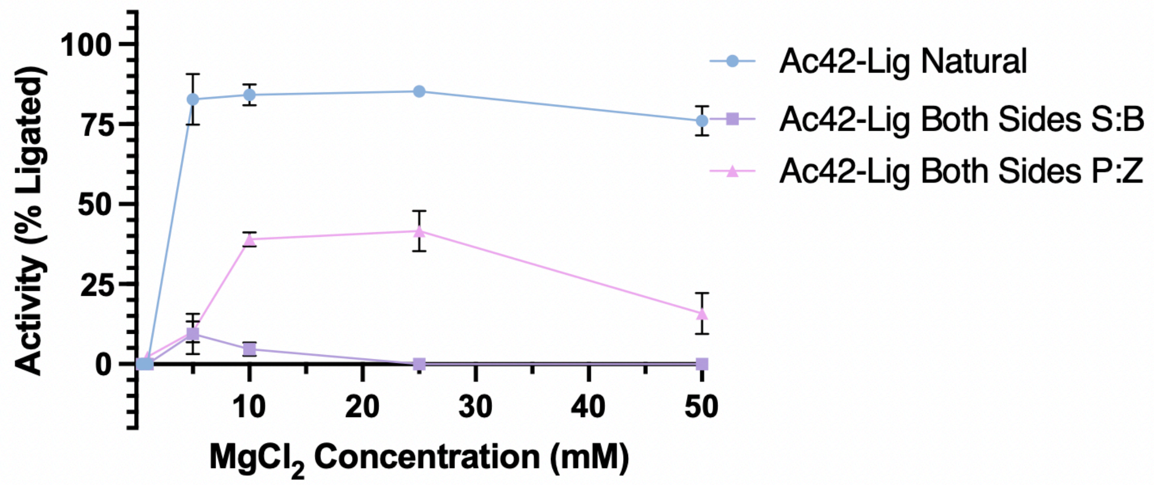

Supplementary 8

DNA Ligase activity of Pmar-LigP and Ac42-Lig with hachimoji substrates and varying manganese ion concentration. Assays were run with standard conditions except Mn concentration varies between 0.1 mM to 5 mM. Error bars are not visible for data points with standard deviation less than 3.0 %.

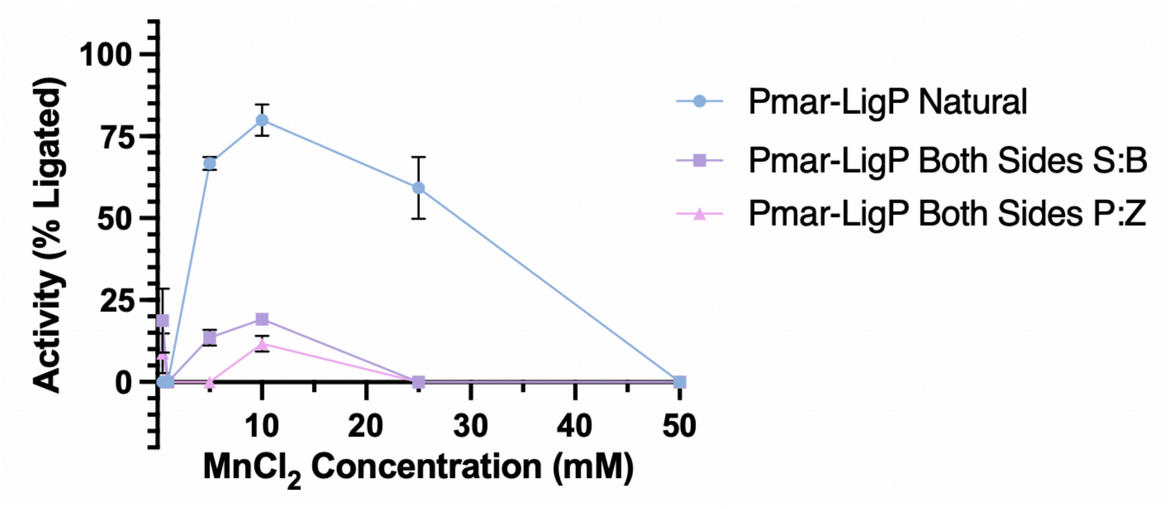


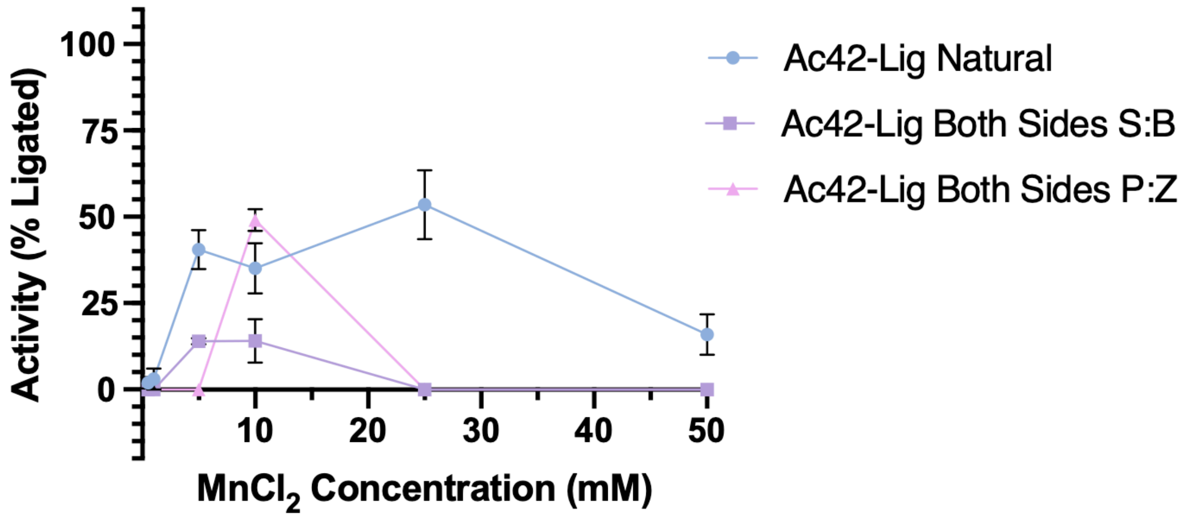

#### Supplementary 9

DNA Ligase activity of Pmar-LigP and Ac42-Lig with hachimoji substrates and extended incubation. Assays were run with standard conditions except incubation time varied between 10 minutes to 1440 minutes. Error bars are not visible for data points with standard deviation less than 3.0 %.

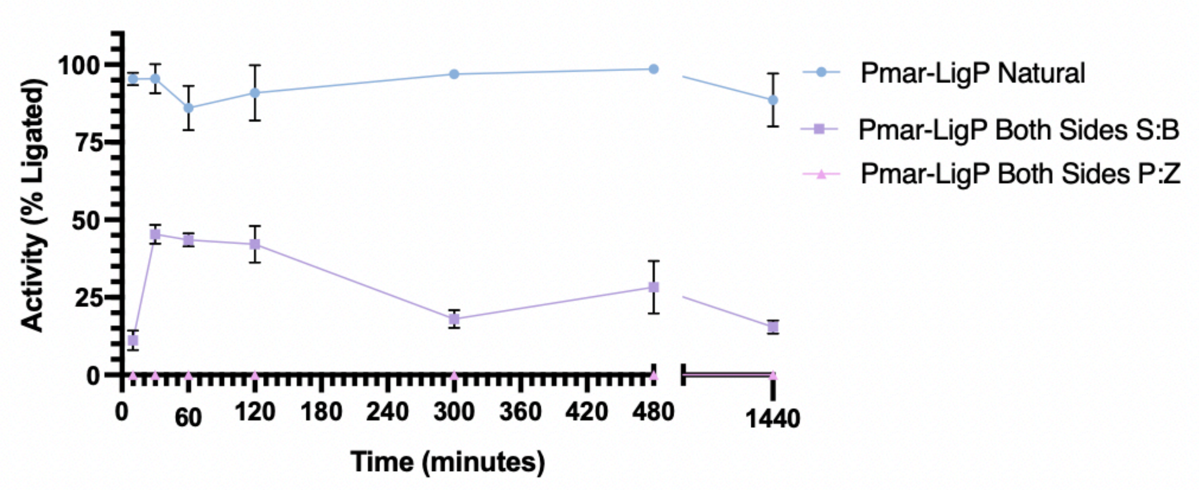


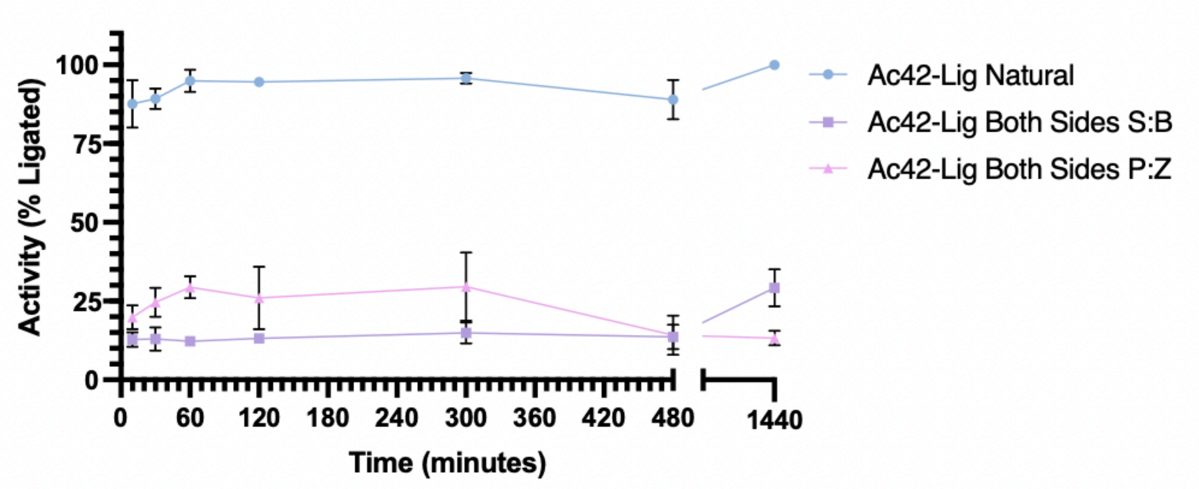


#### Supplementary 10

Duplexes for substrate containing mismatches of canonical or AEGIS bases. The upper strand is written 5' → 3', the complement is presented in the 3'→ 5' orientation for clarity. The individual oligonucleotide sequences and combinations annealed to generate these are provided in Supplementary section 2

**Natural Mismatches**

| DNA Substrate | Oligonucleotides | Sequence 5'-3' (top), 3'-5' (bottom) |
|---|---|---|

| | (Acceptor, Donor, Complement) | |
|---|---|---|
| Nicked | NL1, NL2, NL3 | AGGCCATGGCTGATATCGCA TAGGCATTCGAGCTCCGTCG<br>TCCGGTACCGACTATAGCGT ATCCGTAAGCTCGAGGCAGC |
| 3' A:C | NL1, NL2, NL10 | AGGCCATGGCTGATATCGCA TAGGCATTCGAGCTCCGTCG<br>TCCGGTACCGACTATAGCGC ATCCGTAAGCTCGAGGCAGC<br>X |
| 3' A:G | NL1, NL2, NL14 | AGGCCATGGCTGATATCGCA TAGGCATTCGAGCTCCGTCG<br>TCCGGTACCGACTATAGCGG ATCCGTAAGCTCGAGGCAGC<br>X |
| 5' C:A | NL1, NL2, NL15 | AGGCCATGGCTGATATCGCA CAGGCATTCGAGCTCCGTCG<br>TCCGGTACCGACTATAGCGT ATCCGTAAGCTCGAGGCAGC<br>X |
| 5' G:A | NL1, NL2, NL16 | AGGCCATGGCTGATATCGCA GAGGCATTCGAGCTCCGTCG<br>TCCGGTACCGACTATAGCGT ATCCGTAAGCTCGAGGCAGC<br>X |

#### AEGIS nick- single mismatch

| DNA Substrate | Oligonucleotides (Acceptor, Donor, Complement) | Sequence 5'-3' (top), 3'-5' (bottom) |
|---|---|---|
| 5' B:A Nick Mismatch | NL1, SB2, NL3 | AGGCCATGGCTGATATCGCA BAGGCATTCGAGCT<br>TCCGGTACCGACTATAGCGT ATCCGTAAGCTCGAGGCAGC<br>X |
| 5' S:A Nick Mismatch | NL1, UB-Ni6, NL3 | AGGCCATGGCTGATATCGCA SAGGCATTCGAGCT<br>TCCGGTACCGACTATAGCGT ATCCGTAAGCTCGAGGCAGC<br>X |
| 5' P:A Nick Mismatch | NL1, UB-Ni2, NL3 | AGGCCATGGCTGATATCGCA PAGGCATTCGAGCT<br>TCCGGTACCGACTATAGCGT ATCCGTAAGCTCGAGGCAGC<br>X |
| 5' Z:A Nick Mismatch | NL1, UB-Ni4, NL3 | AGGCCATGGCTGATATCGCA ZAGGCATTCGAGCT<br>TCCGGTACCGACTATAGCGT ATCCGTAAGCTCGAGGCAGC<br>X |
| 3' B:T Nick Mismatch | UB-Ni14, NL2, NL3 | TGGCTGATATCGCB TAGGCATTCGAGCTCCGTCG<br>TCCGGTACCGACTATAGCGT ATCCGTAAGCTCGAGGCAGC<br>X |
| 3' pS:T Nick Mismatch | UB-Ni9, NL2, NL3 | TGGCTGATATCGCS TAGGCATTCGAGCTCCGTCG<br>TCCGGTACCGACTATAGCGT ATCCGTAAGCTCGAGGCAGC<br>X |
| 3' P:T Nick Mismatch | UB-Ni1, NL2, NL3 | TGGCTGATATCGCP TAGGCATTCGAGCTCCGTCG<br>TCCGGTACCGACTATAGCGT ATCCGTAAGCTCGAGGCAGC<br>X |

#### AEGIS complement- single mismatch

| DNA Substrate | Oligonucleotides (Acceptor, Donor, Complement) | Sequence 5'-3' (top), 3'-5' (bottom) |
|---|---|---|
| **5' T:B Comp Mismatch** | NL1, NL2, MM3 | AGGCCATGGCTGATATCGCA TAGGCATTCGAGCTCCGTCG<br>TCCGGTACCGACTATAGCGT BTCCGTAAGCTCGAGGCAGC<br>X |
| **5' T:S Comp Mismatch** | NL1, NL2, MM4 | AGGCCATGGCTGATATCGCA TAGGCATTCGAGCTCCGTCG<br>TCCGGTACCGACTATAGCGT STCCGTAAGCTCGAGGCAGC<br>X |
| **5' T:P Comp Mismatch** | NL1, NL2, MM5 | AGGCCATGGCTGATATCGCA TAGGCATTCGAGCTCCGTCG<br>TCCGGTACCGACTATAGCGT PTCCGTAAGCTCGAGGCAGC<br>X |
| **5' T:Z Comp Mismatch** | NL1, NL2, MM6 | AGGCCATGGCTGATATCGCA TAGGCATTCGAGCTCCGTCG<br>TCCGGTACCGACTATAGCGT ZTCCGTAAGCTCGAGGCAGC<br>X |
| **3' A:B Comp Mismatch** | NL1, NL2, MM7 | AGGCCATGGCTGATATCGCA TAGGCATTCGAGCTCCGTCG<br>TCCGGTACCGACTATAGCGB ATCCGTAAGCTCGAGGCAGC<br>X |
| **3' A:S Comp Mismatch** | NL, NL2, MM8 | AGGCCATGGCTGATATCGCA TAGGCATTCGAGCTCCGTCG<br>TCCGGTACCGACTATAGCGS ATCCGTAAGCTCGAGGCAGC<br>X |
| **3' A:P Comp Mismatch** | NL, NL2, MM9 | AGGCCATGGCTGATATCGCA TAGGCATTCGAGCTCCGTCG<br>TCCGGTACCGACTATAGCGP ATCCGTAAGCTCGAGGCAGC<br>X |

#### AEGIS nick- double mismatch

| DNA Substrate | Oligonucleotides (Acceptor, Donor, Complement) | Sequence 5'-3' (top), 3'-5' (bottom) |
|---|---|---|
| **P-P Nick Mismatch** | UB-Ni1, UB-Ni2, NL3 | TGGCTGATATCGCP PAGGCATTCGAGCT<br>TCCGGTACCGACTATAGCGT ATCCGTAAGCTCGAGGCAGC<br>X X |
| **P-Z Nick Mismatch** | UB-Ni1, UB-Ni4, NL3 | TGGCTGATATCGCP ZAGGCATTCGAGCT<br>TCCGGTACCGACTATAGCGT ATCCGTAAGCTCGAGGCAGC<br>X X |
| **P-S Nick Mismatch** | UB-Ni1, UB-Ni6, NL3 | TGGCTGATATCGCP SAGGCATTCGAGCT<br>TCCGGTACCGACTATAGCGT ATCCGTAAGCTCGAGGCAGC<br>X X |
| **P-B Nick Mismatch** | UB-Ni1, SB2, NL3 | TGGCTGATATCGCP BAGGCATTCGAGCT<br>TCCGGTACCGACTATAGCGT ATCCGTAAGCTCGAGGCAGC<br>X X |
| **S-S Nick Mismatch** | UB-Ni9, UB-Ni6, NL3 | TGGCTGATATCGCS SAGGCATTCGAGCT<br>TCCGGTACCGACTATAGCGT ATCCGTAAGCTCGAGGCAGC |
| **S-B Nick Mismatch** | UB-Ni9, SB2, NL3 | TGGCTGATATCGCS BAGGCATTCGAGCT<br>TCCGGTACCGACTATAGCGT ATCCGTAAGCTCGAGGCAGC<br>X X |
| **S-P Nick Mismatch** | UB-Ni9, UB-Ni2, NL3 | TGGCTGATATCGCS PAGGCATTCGAGCT<br>TCCGGTACCGACTATAGCGT ATCCGTAAGCTCGAGGCAGC |

| | | |
|---|---|---|
| | | X X |
| **S-Z Nick Mismatch** | UB-Ni9, UB-Ni4, NL3 | TGGCTGATATCGCS ZAGGCATTCGAGCT<br>TCCGGTACCGACTATAGCGT ATCCGTAAGCTCGAGGCAGC<br>X X |
| **B-B Nick Mismatch** | UB-Ni14, SB2, NL3 | TGGCTGATATCGCB BAGGCATTCGAGCT<br>TCCGGTACCGACTATAGCGT ATCCGTAAGCTCGAGGCAGC<br>X X |
| **B-S Nick Mismatch** | UB-Ni14, UB-Ni6, NL3 | TGGCTGATATCGCB SAGGCATTCGAGCT<br>TCCGGTACCGACTATAGCGT ATCCGTAAGCTCGAGGCAGC<br>X X |
| **B-P Nick Mismatch** | UB-Ni14, UB-Ni2, NL3 | TGGCTGATATCGCB PAGGCATTCGAGCT<br>TCCGGTACCGACTATAGCGT ATCCGTAAGCTCGAGGCAGC<br>X X |
| **B-Z Nick Mismatch** | UB-Ni14, UB-Ni4, NL3 | TGGCTGATATCGCB ZAGGCATTCGAGCT<br>TCCGGTACCGACTATAGCGT ATCCGTAAGCTCGAGGCAGC<br>X X |

#### AEGIS complement- double mismatch

| **DNA Substrate** | **Oligonucleotides (Acceptor, Donor, Complement)** | **Sequence 5'-3' (top), 3'-5' (bottom)** |
|---|---|---|
| **Z-Z Comp Mismatch** | NL1, NL2, UB-Ni3 | AGGCCATGGCTGATATCGCA TAGGCATTCGAGCTCCGTCG<br>ACCGACTATAGCGZ ZTCCGTAAGCTCGA<br>X X |
| **Z-P Comp Mismatch** | NL1, NL2, UB-Ni5 | AGGCCATGGCTGATATCGCA TAGGCATTCGAGCTCCGTCG<br>ACCGACTATAGCGZ PTCCGTAAGCTCGA<br>X X |
| **Z-B Comp Mismatch** | NL1, NL2, UB-Ni7 | AGGCCATGGCTGATATCGCA TAGGCATTCGAGCTCCGTCG<br>ACCGACTATAGCGZ BTCCGTAAGCTCGA<br>X X |
| **Z-S Comp Mismatch** | NL1, NL2, UB-Ni8 | AGGCCATGGCTGATATCGCA TAGGCATTCGAGCTCCGTCG<br>ACCGACTATAGCGZ STCCGTAAGCTCGA<br>X X |
| **B-B Comp Mismatch** | NL1, NL2, UB-Ni10 | AGGCCATGGCTGATATCGCA TAGGCATTCGAGCTCCGTCG<br>ACCGACTATAGCGB BTCCGTAAGCTCGA<br>X X |
| **B-S Comp Mismatch** | NL1, NL2, UB-Ni11 | AGGCCATGGCTGATATCGCA TAGGCATTCGAGCTCCGTCG<br>ACCGACTATAGCGB STCCGTAAGCTCGA<br>X X |
| **B-Z Comp Mismatch** | NL1, NL2, UB-Ni12 | AGGCCATGGCTGATATCGCA TAGGCATTCGAGCTCCGTCG<br>ACCGACTATAGCGB ZTCCGTAAGCTCGA<br>X X |
| **B-P Comp Mismatch** | NL1, NL2, UB-Ni13 | AGGCCATGGCTGATATCGCA TAGGCATTCGAGCTCCGTCG<br>ACCGACTATAGCGB PTCCGTAAGCTCGA<br>X X |
| **S-S Comp Mismatch** | NL1, NL2, UB-Ni15 | AGGCCATGGCTGATATCGCA TAGGCATTCGAGCTCCGTCG<br>ACCGACTATAGCGS STCCGTAAGCTCGA<br>X X |
| **S-B Comp Mismatch** | NL1, NL2, UB-Ni16 | AGGCCATGGCTGATATCGCA TAGGCATTCGAGCTCCGTCG<br>ACCGACTATAGCGS BTCCGTAAGCTCGA |
| **S-Z Comp Mismatch** | NL1, NL2, UB-Ni17 | AGGCCATGGCTGATATCGCA TAGGCATTCGAGCTCCGTCG<br>ACCGACTATAGCGS ZTCCGTAAGCTCGA<br>X X |
| **S-P Comp Mismatch** | NL1, NL2, UB-Ni18 | AGGCCATGGCTGATATCGCA TAGGCATTCGAGCTCCGTCG |

| | | |
|---|---|---|
| | | ACCGACTATAGCGS PTCCGTAAGCTCGA<br>X X |

#### Consecutive AEGIS separate termini

| DNA Substrate | Oligonucleotides (Acceptor, Donor, Complement) | Sequence 5'-3' (top), 3'-5' (bottom) |
|---|---|---|
| **3' nick SBBC mismatch** | SB14, NL2, NL3 | TGGCTGATATSBBC TAGGCATTCGAGCTCCGTCG<br>TCCGGTACCGACTATAGCGT ATCCGTAAGCTCGAGGCAGC<br>XXXX |
| **5' nick BSS mismatch** | NL1, SB16, NL3 | GGCCATGGCTGATATCGCA BSSGCATTCGAGCT<br>TCCGGTACCGACTATAGCGT ATCCGTAAGCTCGAGGCAGC<br>XXX |
| **3' nick PPP mismatch** | UB14, NL2, NL3 | TGGCTGATATCPPP TAGGCATTCGAGCTCCGTCG<br>TCCGGTACCGACTATAGCGT ATCCGTAAGCTCGAGGCAGC<br>XXX |
| **5' nick ZZZ mismatch** | NL1, UB16, NL3 | GGCCATGGCTGATATCGCA ZZZGCATTCGAGCT<br>TCCGGTACCGACTATAGCGT ATCCGTAAGCTCGAGGCAGC<br>XXX |
| **3' complement GSSB mismatch** | NL1, NL2, SB15 | AGGCCATGGCTGATATCGCA TAGGCATTCGAGCTCCGTCG<br>ACCGACTATABSSG ATCCGTAAGCTCGA<br>XXXX |
| **5' complement BBS mismatch** | NL1, NL2, SB17 | AGGCCATGGCTGATATCGCA TAGGCATTCGAGCTCCGTCG<br>ACCGACTATAGCGT SBBCGTAAGCTCGA<br>XXX |
| **3' complement ZZZ mismatch** | NL1, NL2, UB15 | AGGCCATGGCTGATATCGCA TAGGCATTCGAGCTCCGTCG<br>ACCGACTATAGZZZ ATCCGTAAGCTCGA<br>XXX |
| **5' complement PPP mismatch** | NL1, NL2, UB17 | AGGCCATGGCTGATATCGCA TAGGCATTCGAGCTCCGTCG<br>ACCGACTATAGCGT PPPCGTAAGCTCGA<br>XXX |

#### Consecutive AEGIS both termini

| DNA Substrate | Oligonucleotides (Acceptor, Donor, Complement) | Sequence 5'-3' (top), 3'-5' (bottom) |
|---|---|---|
| **Nick SSBC-BBS mismatch** | SB14, SB16, NL3 | TGGCTGATATSBBC BSSGCATTCGAGCT<br>TCCGGTACCGACTATAGCGT ATCCGTAAGCTCGAGGCAGC<br>XXXX XXX |
| **Nick SSBC-ZZZ mismatch** | SB14, UB16, NL3 | TGGCTGATATSBBC ZZZGCATTCGAGCT<br>TCCGGTACCGACTATAGCGT ATCCGTAAGCTCGAGGCAGC<br>XXXX XXX |
| **Nick PPP-BSS mismatch** | UB14, SB16, NL3 | TGGCTGATATCPPP BSSGCATTCGAGCT<br>TCCGGTACCGACTATAGCGT ATCCGTAAGCTCGAGGCAGC<br>XXX XXX |

| | | |
|---|---|---|
| **Nick PPP-ZZZ mismatch** | UB14, UB16, NL3 | TGGCTGATATCPPP ZZZGCATTCGAGCT<br>TCCGGTACCGACTATAGCGT ATCCGTAAGCTCGAGGCAGC<br>XXX XXX |
| **Complement BBS-GSSB mismatch** | NL1, NL2, SB18 | AGGCCATGGCTGATATCGCA TAGGCATTCGAGCTCCGTCG<br>ACCGACTATABSSG SBBCGTAAGCTCGA<br>XXXX XXX |
| **Complement PPP-ZZZ mismatch** | NL1, NL2, UB18 | AGGCCATGGCTGATATCGCA TAGGCATTCGAGCTCCGTCG<br>ACCGACTATAGZZZ PPPCGTAAGCTCGA<br>XXX XXX |

### Supplementary 11

Structures of DNA ligases bound to natural DNA indicating the position of AEGIS bases used in this study. A) Pmar-LigP bound to natural DNA indicating the interacting side chains with the four nucleotides before the nick (3' acceptor side) and three nucleotides after the nick (5' donor side). Respective zoom regions show electrostatic surface of protein and side-chains of interacting protein residues. Note that no amino acids contact the nucleobase portion of the DNA. For details see (Williamson and Leiros 2019). B) Structures of other ligases in this study with the substituted region shown in red.

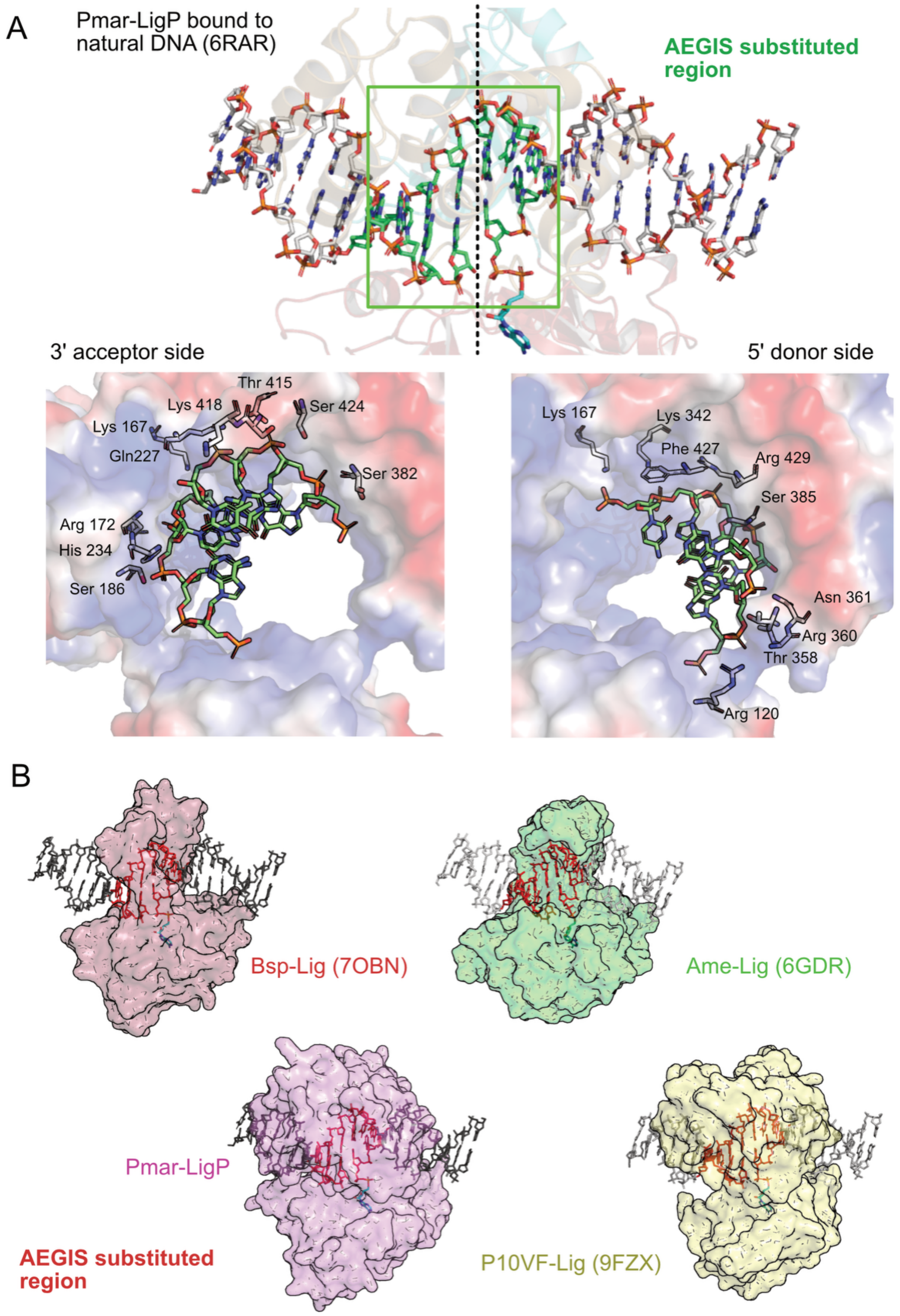

#### Supplementary 12

Mis pairs between natural and AEGIS bases tested in ligation fidelity assays.

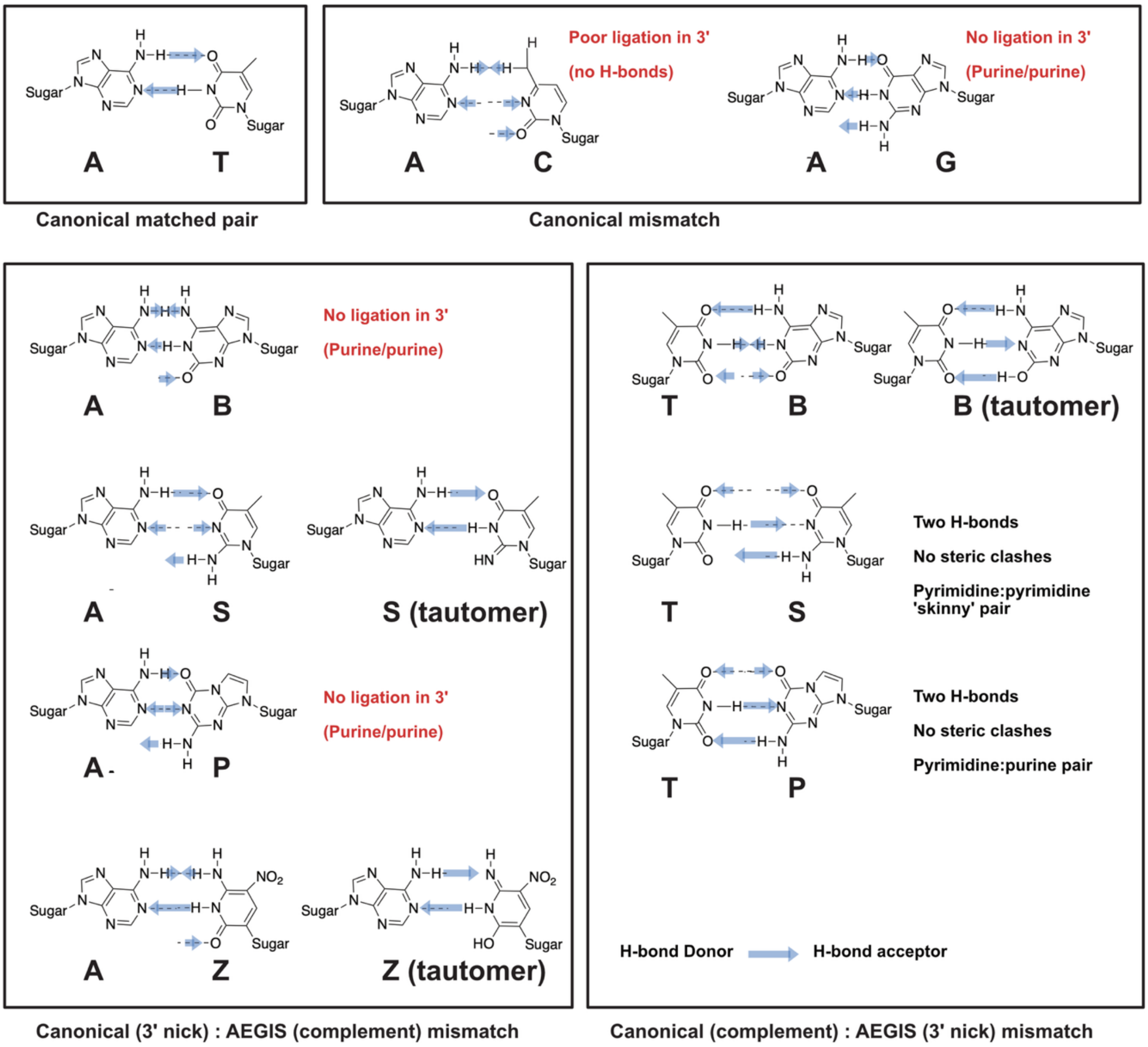
